# Multi-scale structure of chromatin condensates rationalizes phase separation and material properties

**DOI:** 10.1101/2025.01.17.633609

**Authors:** Huabin Zhou, Jan Huertas, M. Julia Maristany, Kieran Russell, June Ho Hwang, Run-wen Yao, Joshua Hutchings, Momoko Shiozaki, Xiaowei Zhao, Lynda K. Doolittle, Bryan A. Gibson, Margot Riggi, Jorge R. Espinosa, Zhiheng Yu, Elizabeth Villa, Rosana Collepardo-Guevara, Michael K. Rosen

**Affiliations:** Department of Biophysics, Howard Hughes Medical Institute, UT Southwestern Medical Center, Dallas, TX, USA; Yusuf Hamied Department of Chemistry, University of Cambridge, Cambridge CB2 1EW, UK; Department of Genetics, University of Cambridge, Cambridge CB2 3EH, UK; Maxwell Centre, Cavendish Laboratory, Department of Physics, University of Cambridge, Cambridge CB3 0HE, U; School of Biological Sciences, University of California San Diego, La Jolla, CA, USA; Janelia Research Campus, Howard Hughes Medical Institute, Ashburn, VA, USA; Howard Hughes Medical Institute, University of California San Diego, La Jolla, CA, USA; Max Planck Institute for Biochemistry, Martinsried/Munich D-82152, Germany; Marine Biological Laboratory Chromatin Collaborative, Marine Biological Laboratory, Woods Hole, MA, USA

## Abstract

Biomolecular condensates, compartments that concentrate molecules without surrounding membranes, are integral to numerous cellular processes. The structure and interaction networks of molecules within condensates remain poorly understood. Using cryo-electron tomography and molecular dynamics simulations we elucidated the structure of phase separated chromatin condensates across scales, from individual amino acids to network architecture. We found that internucleosomal DNA linker length controls nucleosome arrangement and histone tail interactions, shaping the structure of individual chromatin molecules both within and outside condensates. This structural modulation determines the balance between intra- and intermolecular interactions, which in turn governs the molecular network, thermodynamic stability, and material properties of chromatin condensates. Mammalian nuclei contain dense clusters of nucleosomes whose non-random organization is mirrored by the reconstituted condensates. Our work explains how the structure of individual chromatin molecules ultimately determines physical properties of chromatin condensates, with implications for cellular chromatin organization.

## Introduction

Cells are organized across a wide range of length scales, from nanometer-scale structures of individual molecules to micrometer-scale subcellular compartments formed by assembly of these components. Biomolecular condensates, which concentrate molecules into discrete foci without surrounding membranes, exemplify how nanoscale interactions can produce micron scale biological organization. Many condensates appear to form through assembly and concomitant phase separation of multivalent macromolecules (1–3). The structures of individual molecules and of their multivalency- enabled interaction networks, plus the dynamics of both entities, strongly influence the macroscopic physical properties of condensates (4–6). Physical properties, along with solution environment and composition define the biochemical and cellular functions of condensates (7–16). These features of condensates are altered in diseases and have been suggested as therapeutic targets (17, 18). Thus, an important goal in biology and human health is to understand the relationships between molecular structure/dynamics, network architecture/dynamics and function of condensates.

Molecules within condensates are often conformationally flexible and interact weakly and transiently, producing complex distributions of interconverting states. NMR spectroscopy has revealed structural ensembles and residue-specific interactions of intrinsically disordered regions of proteins (IDRs) within condensates (19–21). But these studies have not strongly informed on the higher order organization of these elements within condensates. On micrometer length scales, cryo-electron tomography has been used to characterize the morphology of condensates (22–29), but has not connected meso-scale features to molecular interactions that form the compartments. Computational work has sought to bridge molecular interactions to the network architecture of intrinsically disordered protein (IDP) condensates, providing insight into viscoelasticity and surface tension (5, 30–32). But these models are difficult to test experimentally due to the challenge of directly visualizing molecules and interactions inside condensates. Because of these complexities, we still lack a complete understanding from molecular structure and interactions to meso-scale organization and dynamics for any condensate.

Phase separation has been hypothesized to play an important role in the organization of eukaryotic chromatin, enabling the compaction necessary for storage in the nucleus while retaining the dynamics necessary for function (33–43). Chromatin fibers consist of large arrays of nucleosomes, structural units containing ∼147 base pairs of DNA wrapped around an octamer of histone proteins (44), connected by DNA linkers of diverse lengths. Nucleosome arrays compact along their lengths and interact with one another (45), creating higher order cellular structures ranging from 10-100 nm clusters (46–48) to 100-300 nm domains (49, 50). The compaction and dynamics of nucleosomes are highly regulated, and play important roles in nuclear processes including transcription (51–53), DNA replication and DNA repair (54). Interactions between nucleosome arrays can drive salt-dependent phase separation in vitro in a manner dependent on the disordered tails of the histone proteins (33, 35, 55–60). Phase separation can also be modulated by the length of inter-nucleosome DNA linkers, as the drive to phase separate shows an oscillatory pattern as linkers are extended, stronger for 10N+5 base pairs (integer N) and weaker for 10N base pairs (33). This process produces substantial nucleosome compaction while retaining dynamics, and is regulated similarly to chromatin in cells (33, 34). These behaviors, along with genomic data and modeling, suggest that phase separation of the chromatin polymer plays a role in genome organization in vivo (33, 35, 57–61).

A variety of studies have examined the structure of native chromatin fibers using different modalities of light and electron microscopy (46, 49, 62–66), as well as chemical crosslinking (67, 68). These have revealed that chromatin inside cells/nuclei consists mostly of irregular conformations, although small segments of ordered, stacked groups of nucleosomes have also been observed (62, 65, 69, 70). Most of these studies focused on the conformations of individual fibers, rather than the structural basis of interactions between them. However, in one study pairwise inter-nucleosome interactions revealed a wide distribution of packing geometries (65). The relationship between the conformations, interactions and networking of fibers to the density and material properties of chromatin in cells is not known.

Here we used a combination of cryo-electron tomography (cryo-ET), computer simulations and light microscopy to understand mechanisms of phase separation and physical properties of synthetic and native chromatin. We found that nucleosome linker length controls salt-dependent conformations and histone tail interactions of chromatin fibers. These properties dictate the strength and number of interactions between fibers in condensates, which then define the network structure of the condensates. These physical features explain the driving force for chromatin phase separation as well as viscoelastic properties of chromatin condensates. Dense, 100-300 nm clusters of native chromatin in situ show a similar, non-random distribution of inter-nucleosome contacts as one class of synthetic chromatin. Together, our studies provide an understanding of chromatin phase separation from nanometers to micrometers, and suggest that certain reconstituted chromatin condensates may functionally mimic important features of native chromatin.

## Results

### Distinct Phase Separation Propensity and Dynamics of 25 bp and 30 bp Chromatin

We generated dodecameric nucleosome arrays based on the Widom 601 positioning sequence (71, 72), with either 25 bp or 30 bp internucleosomal DNA linkers (25 bp or 30 bp chromatin, respectively). As previously reported (33), we found that the 25 bp chromatin phase separates at lower concentration and at lower ionic strength than 30 bp chromatin (Fig. 1A). In partial droplet fluorescence recovery after photobleaching (FRAP) experiments 25 bp chromatin recovers appreciably more slowly than 30 bp chromatin (Fig. 1B). Further, 25 bp chromatin droplets also fuse more slowly than 30 bp chromatin droplets (Fig. 1C), with longer relaxation times suggesting higher viscoelasticity and/or lower surface tension (Fig. 1D). We combined cryo-ET and multiscale molecular dynamics simulations to understand these differences in drive to phase separate and material properties (Fig. 1E).

**Figure 1.**
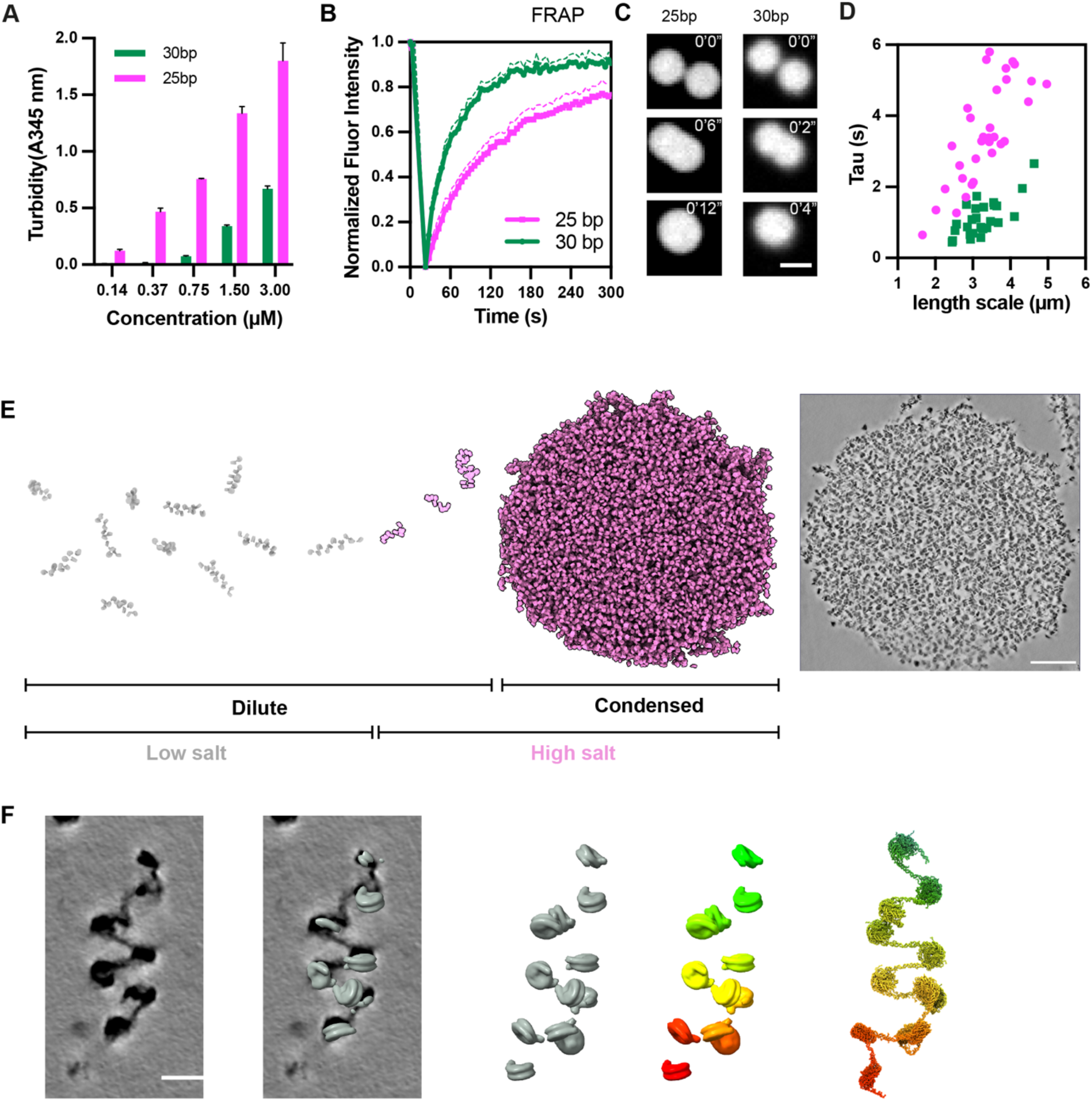
Distinct phase separation propensities and dynamics of 25 bp and 30 bp chromatin. **(A)** Measurements of 25 bp and 30 bp chromatin turbidity with absorption at 345 nm at indicated nucleosome concentrations. Error bars indicate ± standard deviation of three measurements. **(B)**Fluorescence Recovery After Photobleaching (FRAP) results for 25 bp and 30 bp chromatin. Error bars indicate ± standard deviation of 20 measurements. **(C)**Time-lapse imagery of droplet fusion events in 25 bp and 30 bp chromatin, with a scale bar representing 3 µm to show relative droplet sizes and fusion rates. **(D)**Measurements of the timescale of fusion for 25 bp and 30 bp chromatin droplets. **(E)**Schematic illustration of the compaction and phase separation of chromatin fibers as salt concentration increases, as visualized by cryo-ET. **(F)** Schematic illustration of the computational approach used to reconstruct histone tails and increase the resolution of a single chromatin array visualized by cryo-ET in low salt conditions. From left to right the images show: chromatin density in a denoised cryo-ET tomogram; nucleosomes placed by CATM into the density; nucleosome models extracted from the density; nucleosomes colored from red (1) to green (12) according to their linear connections in the fiber; higher resolution fiber computationally reconstructed using molecular dynamics simulations steered according to nucleosome positions in the cryo-ET data. Further simulations restraining nucleosomes to their cryo-ET positions and orientations produce structural ensembles of the histone tails.

### Salt differentially compacts and reorganizes histone tail interactions in 25 bp and 30 bp chromatin

We collected cryo-ET data in low- and high salt conditions (0 and 150 mM potassium acetate, respectively) for solutions of 25 bp and 30 bp chromatin. In low salt, the molecules are monomeric and we could use standard blotting techniques to produce vitrified samples for imaging and analysis (Fig. 1F). In high salt, the chromatin solution phase separates, and we used the Waffle method (73), combining high-pressure freezing (HPF) and cryogenic Focused Ion Beam (cryo-FIB) milling, to produce thin lamellae containing cylindrical slabs sliced out of spherical droplets (Fig. 1E, S1A) (74). For all samples, we collected tilt-series, and then reconstructed and denoised tomograms. In low salt conditions we could readily identify nucleosomes and trace their connectivity in individual chromatin fibers in the tomograms (Fig. 1F, S1B).

Using these procedures we first sought to understand how increasing salt affects the structure of individual nucleosome arrays to promote phase separation. Numerous biophysical studies have shown that mono- and di-valent salts cause compaction and oligomerization of chromatin fibers (75–81), although most studies have not revealed the underlying structural details (but see (82) and (83) for recent exceptions). To analyze the chromatin structures we measured the distance between the centers of mass, D, and the dihedral angle between planes, para, for all pairs of nucleosomes N and N+2 in each array (where N represents sequential position). We also measured the dihedral angle between the planes of adjacent (N, N+1) nucleosomes, α (Fig. 2A-D). To assist in visualizing these geometric features, in each system/condition we reconstructed an average di-nucleosome, and then combined two such pairs to build an average trinucleosome model (fig. S1C), shown in Figures 2E and F.

**Figure 2.**
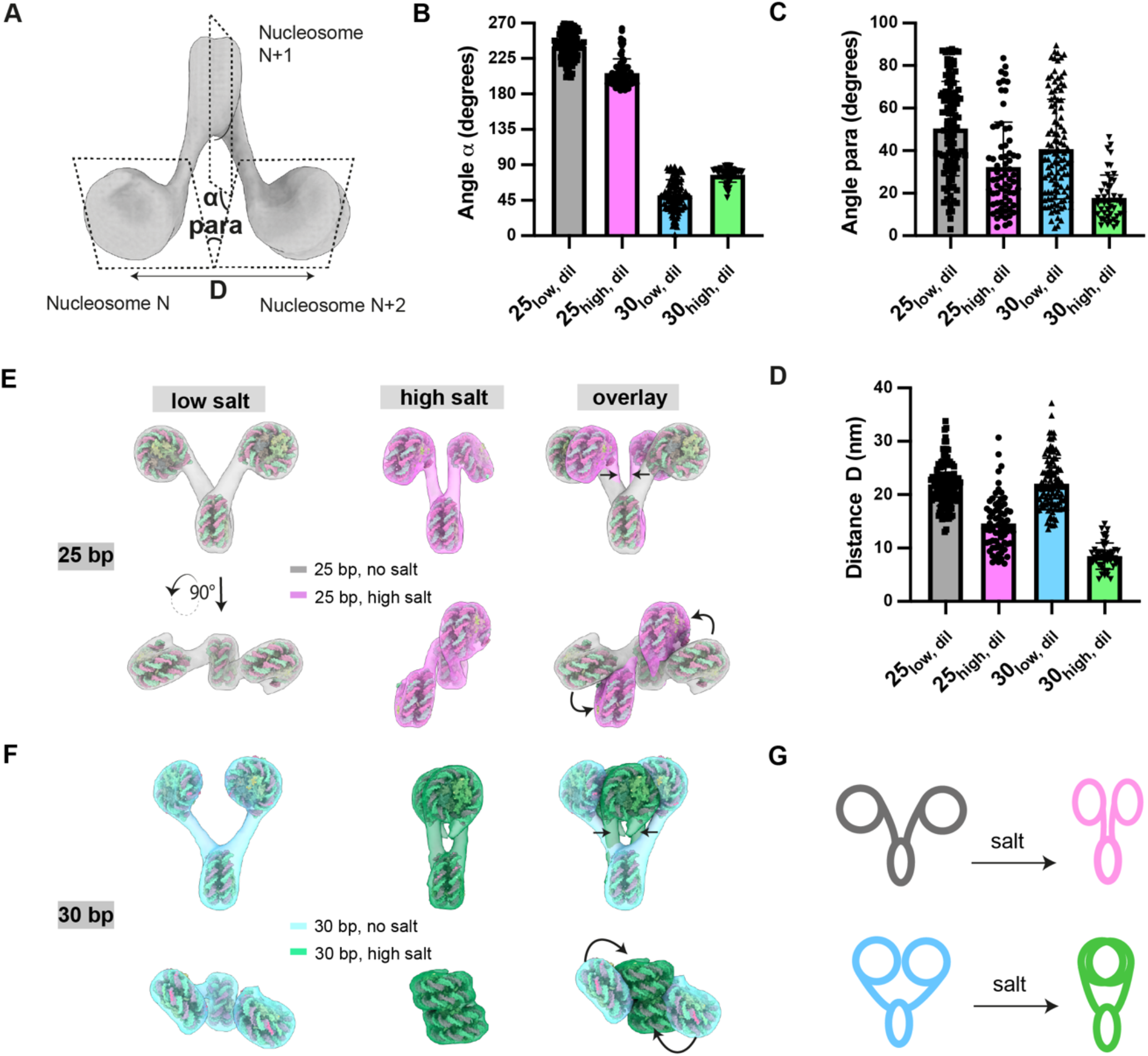
Linker length tunes nucleosome phasing and histone tail interactions, producing different structures in response to salt in the dilute phase. **(A)** Schematic illustration of nucleosome features measured in this study. **(B-D)** Statistical characterization of the dihedral angle between consecutive nucleosomes (α), the dihedral angle between nucleosomes N and N+2 (para), and the distance between nucleosomes N and N+2 (D). Chromatin with 25 bp spacing was colored gray under low-salt dilute conditions (25low, dil) and light magenta under high-salt dilute conditions (25high, dil). Similarly, chromatin with 30 bp spacing was colored light blue under low-salt dilute conditions (30low, dil) and light green under high-salt dilute conditions (30high, dil). **(E-F)** Representative structures of chromatin with 25 bp (a) and 30 bp (b) linker lengths at low salt concentration in the dilute phase (left) and at high salt concentration in the dilute phase (middle). The right panels show overlays of the respective structures to highlight differences in condensation behavior under varied ionic conditions. **(G)** Schematic representation of the conformational changes induced by salt in 25 bp and 30 bp chromatin.

In low salt, the 25 bp and 30 bp chromatin have similarly large values of D (Fig. 2D), indicating that semi-adjacent N and N+2 nucleosomes are splayed apart from one other; para values are ∼45 ° in both cases. The average α angle for the two arrays differs by nearly 180° (α ∼ 230° and 50°, for 25 bp and 30 bp, respectively, Fig. 2B), reflecting the difference in phasing of successive nucleosomes imparted by the 10 bp/turn helical pitch of the linker DNA. Because of this difference, in 25 bp chromatin the N and N+2 nucleosomes face outward, away from each other (Fig. 2E, left), whereas in 30 bp chromatin they face inward, toward each other (Fig. 2F, left).

Increased salt reduces long-range electrostatic repulsion in the chromatin DNA (79), leading to rearrangements and compaction of both array types (fig. S1B). In both cases, D decreases appreciably (Fig. 2D), as does para (Fig. 2C). In the 25 bp chromatin, nucleosomes N and N+2 remain unstacked with both faces exposed to solvent (Fig. 2E), and become roughly parallel to the intervening N+1 nucleosome as shown by the decrease in α (to ∼ 200°, Fig. 2B, E). The changes to D and para are larger in the 30 bp chromatin, due to face-to-face stacking of the N and N+2 nucleosomes, which become oriented approximately perpendicular to the N+1 nucleosome (Fig. 2F) as indicated by the large increase in α (to ∼80°, Fig. 2B). Similar but smaller decreases in D and para were recently noted in compaction of 40 bp tetra-nucleosome arrays between 5 and 50 mM NaCl (83). Notably, computational studies have shown that face-to-face and face-to-side contacts are the most energetically favorable nucleosome packing geometries, followed distantly by side-to-side (84, 85). These changes result in greater compaction of 30 bp chromatin than 25 bp chromatin, as reflected by the smaller radius of gyration of the former (fig. S1D), due to intramolecular stacking of nucleosomes. When stacking is repeated across the 30 bp arrays, it produces two-start helices similar to, but appreciably less regular than, those observed in crystal structures (86) and single-particle cryo-EM reconstructions (87) of arrays with linkers of 20, 30 or 40 base pairs (fig. S1E).

To gain insight into the histone tail interactions driving chromatin compaction with increasing salt, we combined experimental cryo-ET data with computational simulations using a GPU-accelerated version of our chemically specific coarse-grained chromatin model (88), which represents chromatin at amino acid and nucleotide resolution (Fig. 3A–D). We simulated a series of 12-nucleosome arrays observed in the cryo-ET tomograms, restraining the nucleosome geometric centers to their experimental positions in low (Fig. 3A–B) and high salt (Fig. 3C–D). The simulations generate structural ensembles of the chromatin-bound histone tails, revealing the probabilities of different histone-tail conformational states. We analyzed the intra-nucleosome and internucleosome contacts (fig. S2A) mediated by histone tails by quantifying the number of amino acid pairs and amino acid– DNA pairs in contact during the simulations. This analysis identifies the most frequent interactions within nucleosome pairs arranged based on the cryo-ET data. However, interaction energy is influenced not only by proximity but also by the chemical nature of the interacting pairs (e.g., lysine to phosphate versus alanine to phosphate). Nevertheless, our analyses show that the most frequent tail-mediated inter-nucleosome interactions arise from energetically favorable pairs, specifically lysines and arginines interacting electrostatically with negatively charged histone residues and DNA (fig. S2B).

**Figure 3.**
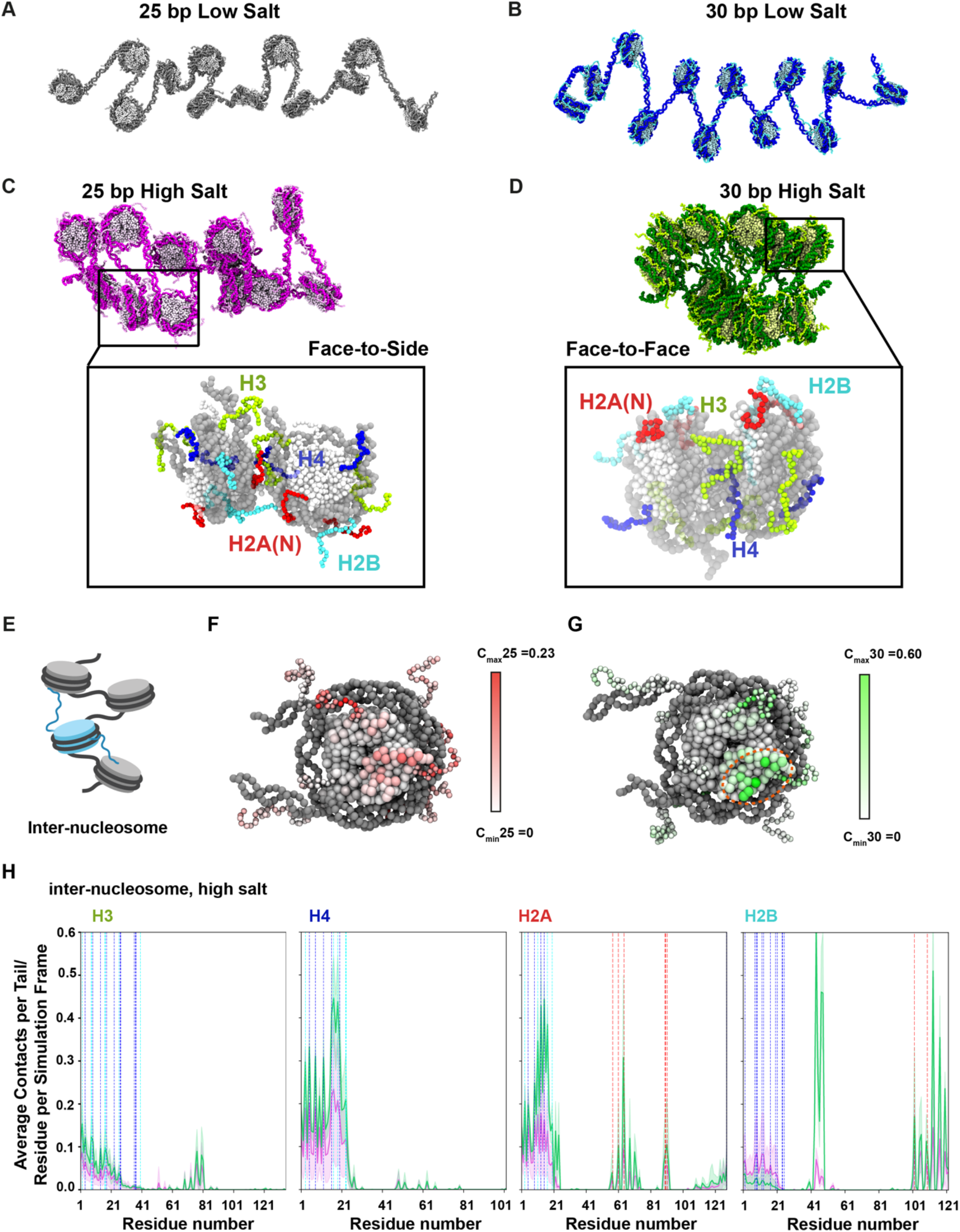
Computer simulations reveal different histone tail interactions during compaction of 25 bp and 30 bp chromatin in the dilute phase. **(A)**Schematic representation of a 25 bp chromatin array derived from cryo-ET and computationally extended to the high resolution chemical-specific model in the dilute phase in low salt. DNA beads are shown in dark grey and histone protein beads in light grey. **(B)**As in A but for a 30 bp chromatin. DNA beads are shown in dark blue and histone protein beads in light blue. **(C)**As in (A) but for 25-bp chromatin in high-salt. DNA beads are shown in dark pink and histone protein beads in light pink. The inset expands the image of one pair of nucleosomes interacting face-to–side via their histone tails. To aid visualization, all other nucleosomes have been deleted, the histone core beads are shown in white, and DNA beads in light grey. Histone tail beads are colored as follows: H3 in green, H4 in blue, H2A-N terminal in red, and H2B in cyan. **(D)**As in (A) but for 30-bp chromatin in high-salt. DNA beads are shown in dark green and histone protein beads in light green. The inset expands the image of one pair of nucleosomes interacting face-to–face via their histone tails. The image in the inset is colored as in (C). **(E)**Schematic representation of inter-nucleosome tail interactions. **(F-G)** Illustration of representative nucleosome structures, showing the histone core as larger beads and the histone tails as smaller beads. Residues are color-coded to represent contact frequency, defined as the total number of inter-nucleosome contacts made by each amino acid in histones (H3, H4, H2A(N), and H2B) of one nucleosome with DNA (nucleosomal and linker) and histones (core and tails) of neighboring nucleosomes. Two scales are used: a finer scale (0–0.23) for 25 bp chromatin (F, left) and a broader scale (0–0.6) for 30 bp chromatin (G). The acid patch in G is emphasized with a dashed orange oval. **(H)** Number of inter-nucleosome contacts mediated by each amino acid in the histone (H3, H4, H2A(N), and H2B) of one nucleosome and the DNA (nucleosomal and linker) and histones (core and tails) of a neighboring nucleosome. Data for 25 bp fibers is shown in pink and for 30 bp fibers in green. The blue, cyan and orange vertical lines show the positions of lysines and arginines, acidic patch residues, respectively. The shading indicates the standard deviation from the mean.

The simulations show that in low salt, where DNA–DNA electrostatic repulsion drives nucleosomes apart (Fig. 3A–B), the histone tails in both chromatin types make almost exclusively intra-nucleosome interactions with DNA (fig. S2C); inter nucleosome contacts are negligible (fig. S2D), as observed in previous simulations (89). In high salt, the tails partially release from their own nucleosomes (fig. S2C) and shift to contact the DNA and histones of neighboring nucleosomes (Fig. 3C–E and fig. S2C). The nature of these interactions is governed by the conformation of the fiber, and also by the position from which each tail emerges from the nucleosome core: H4 and H2A(N) emerge from the nucleosome face, whereas H3 and H2B pass between the two gyres of DNA and extend from the nucleosome edge (Fig. 3C-D and fig. S2E).

In high salt, the 25 bp arrays adopt a heterogeneous distribution of conformations with nucleosomes displaying a broad range of distances and orientations relative to their neighbors (Fig. 2B-D and S1B). This diversity in inter-nucleosome configurations coincides with highly heterogeneous interactions involving all histone tails (Fig. 3C, E, F and H). This heterogeneity is evident in the large standard deviations from the mean in the number of contacts per residue (Fig. 3H), indicating that the contributions of different tails are comparable within these deviations. The H3 tail, projecting from the nucleosome side, is ideally positioned for face-to-side and side-to-side contacts between nucleosomes, while the H4 and H2A(N) tails, projecting from the nucleosome face, facilitate face-to-side interactions and, less frequently, face-to-face contacts (Fig. 3C). Consistent with the diverse tail-behaviors of the 25 bp chromatin, simulations indicate that the free energy landscape of a 25-bp tetra-nucleosome is characterized by multiple competing low-lying minima (90).

The case is different for the 30 bp arrays, where intramolecular face-to-face nucleosome stacking dominates. The prevalence of this geometry decreases the heterogeneity of tail interactions (strongly reducing their standard deviations from the mean in Fig. 3E), and amplifies the contributions of the H4 and H2A(N) tails, whose positions facilitate stabilization of face-to-face stacking (Fig. 3D–E and S2E). The H3 and H2B tails are less involved due to their peripheral locations. Because of the greater compaction of 30 bp arrays, the H4, H2A(N), and H3 tails make appreciably more inter-nucleosome contacts than in 25 bp chromatin (Fig. 3E). The region around H4K16 (including K16, R17, and R19) establishes the highest number of inter-nucleosome contacts (Fig. 3E and S3B), in agreement with the effect of H4K16 acetylation or mutation of this so-called basic patch in triggering chromatin decompaction (91–93). Residues in and adjacent to the H2A and H2B acidic patch (H2A residues Y57, E61, E64, L65, N68, R71, D72, N89-E92; H2B residues E102, E110 plus L42-D47) formed more interactions in 30 bp chromatin compared to 25 bp chromatin (Fig. 3F-H), primarily due to interactions with the H4-tail basic patch (94–97).

As detailed in Supplemental Tables 1-2, the histone tail contact distributions of 25 bp and 30 bp chromatin are consistent with previous biochemical studies on the roles of individual tails on chromatin compaction. Previous simulations performed without experimental restraints have also revealed that the role of specific histone tails in mediating inter-nucleosome interactions is dependent on solution conditions, linker DNA length, and inter-nucleosome orientations (89, 98–101), as we observed here.

Overall, our data and simulations show that increased salt concentration drives compaction of nucleosome arrays and concomitant reorganization of histone tails from almost entirely intra-nucleosome contacts to also include many inter-nucleosome contacts. For 25 bp chromatin, DNA geometry and diverse contacts of the H3, H4 and H2A(N) histone tails produce heterogeneous structures with many nucleosome faces exposed to solvent. In contrast, for 30 bp chromatin, DNA geometry and interactions of histone H4 and H2A(N) tails favor intramolecular stacking of nucleosomes. As described below, these differences in configuration strongly influence the ability of molecules to interact with neighbors, with important consequences on phase separation and condensate material properties.

### 25 bp and 30 bp chromatin form distinct molecular structures in the condensates

To understand how molecular features impact phase separation and condensate properties, we next examined the structure of chromatin in the condensed phase. Using recently developed context-aware template matching (CATM) algorithms (74), we placed and oriented the nucleosomes for both 25 bp and 30 bp chromatin condensates (Fig. 4A, E). Nearly random orientation distributions with respect to the imaging axis in both cases indicate that sample integrity was preserved and nucleosomes were properly assigned (fig. S3A, (74)). Subtomogram averaging resulted in nucleosome structures with sub-nanometer resolution without discarding any particles—8 Å for 25 bp chromatin, based on 109,801 particles in 10 tomograms, and 7 Å for 30 bp chromatin, based on 119,577 particles in 10 tomograms (fig. S3B)—again supporting the accuracy of nucleosome positions and orientations.

**Figure 4.**
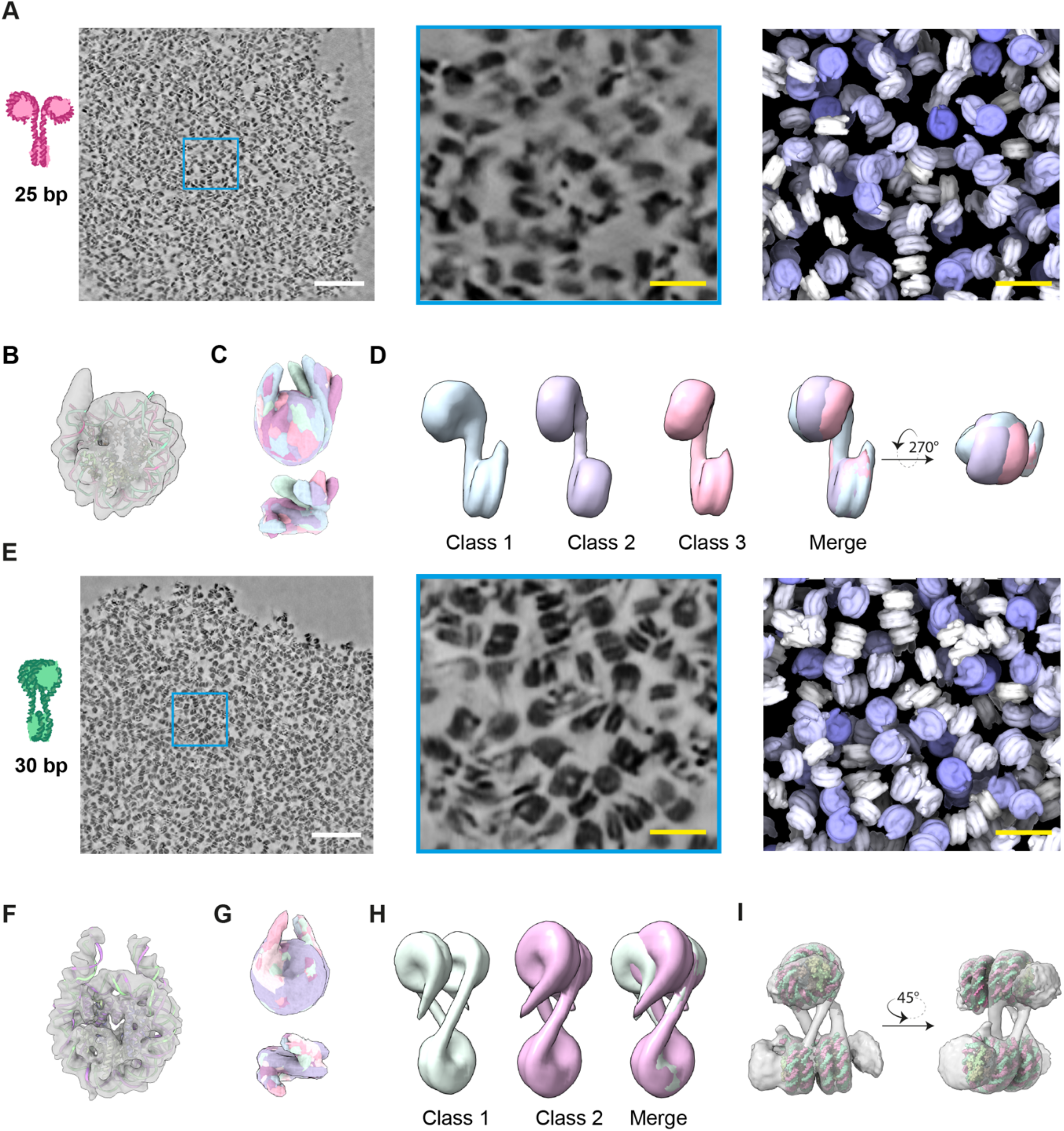
Different molecular structures of 25 bp and 30 bp chromatin the condense phase. **(A)**Tomographic analysis of 25 bp chromatin. Left panel shows a cross-sectional view of a chromatin condensate (scale bar is 100 nm). Center panel magnifies the region within the blue square of the left image (scale bar is 20 nm). Right panel shows nucleosome model assignments from densities in the center panel (scale bar is 20 nm). **(B)**Nucleosome structure from 25 bp chromatin obtained by sub-tomogram averaging 109,801 particles in 10 tomograms, with 8.1 Å resolution. Density shown as a grey surface, fitted model (PDB: 6pwe) is shown in red and green ribbons. **(C)**Classification of mono-nucleosome structures in 25 bp chromatin. Each class is colored differently. **(D)**Classification of di-nucleosome structures within 25 bp chromatin. Each class is colored differently. **(E)**Overview of the 30 bp chromatin structure, analogous to panel A. **(F)**Nucleosome structure from 30 bp chromatin obtained by sub-tomogram averaging 119,577 particles in 10 tomograms, with 7.6 Å resolution. Density shown as a grey surface, fitted model (PDB: 6L4A) is shown in red and green ribbons. **(G)**Classification of mono-nucleosome structures in 30 bp chromatin shown in overlapped density. Each class is colored differently. **(H)**Two classes of tri-nucleosome structures within 30 bp chromatin. The two structures have different degrees of overlap between the nucleosome faces (left and middle images). Each class is colored differently, and an overlay is shown on the right. **(I)**Structural modeling of a tetra-nucleosome in 30 bp chromatin. Density shown as a grey surface, fitted four individual mono-nucleosome models (PDB: 6pwe) are shown in red and green spheres.

During high-resolution refinement, we found that nucleosomes in 25 bp chromatin showed only one linker DNA segment emerging from the core particle (Fig. 4B), suggesting flexibility within the condensed phase. Subclassification yielded a series of nearly identical mono-nucleosome structures, but showed a panel of distinct orientations of the second linker when aligned based on the first (Fig. 4C). These orientations differed in both the DNA crossing angle at the nucleosome dyad (Fig. 4C, top) and also in deviations out of the nucleosome plane (Fig. 4C, bottom), similar to mono-nucleosomes analyzed in high salt (102). In further reconstructions of 25 bp chromatin di-nucleosomes, we found that adjacent nucleosomes can adopt different conformations, with varying twist angles (α) between them (Fig. 4D). The variability in twist angle between classes is comparable to that of di-nucleosome pairs in dilute phase molecules (fig. S3C), indicating that flexibility is not specific to either phase. This flexibility prevented subtomogram averaging of tri-nucleosomes or more complex structures. The flexibility of both individual nucleosomes and nucleosome pairs is likely related to the reported importance of nucleosome breathing and DNA twisting for efficient chromatin phase separation (57, 88).

The 30 bp chromatin behaves quite differently (Fig. 4E). -nucleosomes in 30 bp chromatin condensates could be refined to high resolution with two linker DNA segments clearly observable (Fig. 4F, G). These segments have a crossing angle of 57°, similar to that observed in cryo-EM reconstructions of mono-nucleosomes (Fig. 3F, G, (103, 104)). By increasing the box size used in subtomogram averaging, we could also reconstruct tri-nucleosome (two conformers, Fig. 4H) and tetra-nucleosome structures, the latter with additional partial nucleosome densities surrounding it (Fig. 4I). Both assemblies show pairs of stacked, alternating nucleosomes (N:N+2), oriented approximately orthogonally to the other nucleosome(s). Efforts to refine larger structures with additional nucleosomes were unsuccessful, suggesting that the tetra-nucleosome is the largest well-ordered element in the 30 bp dodecameric arrays within the condensates. This structural heterogeneity beyond a tetra-nucleosome likely reflects the heterogeneity in dynamics observed previously by single molecule FRET (105) and other analytical tools (106).

Overall, within the condensates the 25 bp chromatin adopts more diverse configurations, while 30 bp chromatin is more constrained and stereotypical. This is true at the level of individual nucleosomes and also in larger groupings within a fiber. As in the dilute phase, the 30 bp chromatin makes numerous intramolecular face-to-face stacking interactions, which are not frequently observed in the 25 bp chromatin.

### Balance of intra- and inter-molecular interactions determines condensate stability

To translate molecular features of the 25 bp and 30 bp chromatin to the phase separation behaviors of the two molecules, we next compared the organization of nucleosomes and nucleosome arrays in the two systems on larger scales. First, the radial distribution function, g(r), of nucleosomes in 25 bp chromatin is similar to that of a random distribution at equivalent density, while that of 30 bp chromatin shows peaks indicating pair-wise and higher order face-to-face stacking (Fig. 5A). Similarly, the distribution of angles between the planes of nearest neighbor nucleosomes in 25 bp chromatin deviates only modestly from a random (sinusoidal) distribution (74), with a small excess at low angles (∼parallel geometry) and minor depletion at angles nearer to 90° (perpendicular geometry) (Fig. 5B). In contrast, nucleosomes in the 30 bp condensates show a strong preference for low angles, with a sharp maximum at ∼15 degrees and substantial depletion for angles >∼45 degrees (Fig. 5B). Together, the radial distribution function and angular analyses are consistent with an abundance of face-to-face nucleosome stacking in the 30 bp condensates and more heterogeneous orientations in the 25 bp condensates.

**Figure 5.**
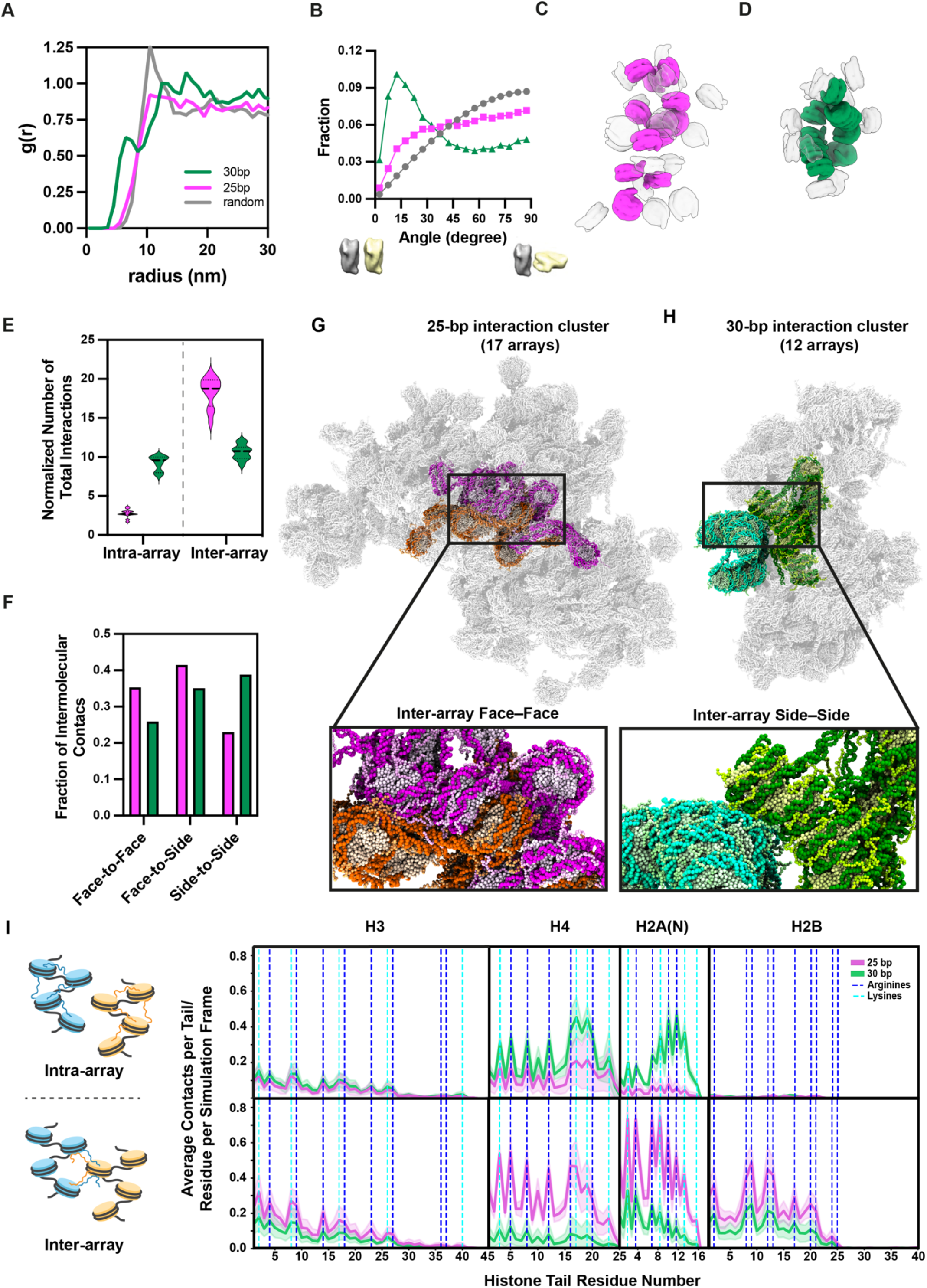
Linker length determines the balance of intraand inter-molecular nucleosome interactions in the condense phase. **(A)**Radial distribution functions, g(r), of 25 bp (magenta) and 30 bp (green) chromatin. 25 bp chromatin has g(r) similar to that of a computed bath of randomly oriented nucleosomes at an equivalent density, with a single maximum at ∼11 nm and a broad distribution at longer lengths. 30 bp chromatin has g(r) with peaks at 6, ∼12, and ∼16 nm. **(B)**Distributions of nearest di-nucleosome orientations for 25 bp (magenta) and 30 bp (green) chromatin. Zero and 90 degrees represent parallel and perpendicular orientations, respectively. **(C, D)** Representative traced individual nucleosome arrays (25 bp magenta in C, 30 bp green in D) and their immediate neighboring nucleosomes (grey) within chromatin condensates. **(E)**Normalized (to 12 nucleosomes) count of intra and inter-nucleosome interactions of 25 bp (magenta) and 30 bp (green) nucleosome arrays within chromatin condensates. **(F)**Fraction of intermolecular contacts between traced arrays and their immediate neighboring nucleosomes with face-to-face, face-to-side and side-to-side geometries (25 bp magenta, 30 bp green). **(G)**Representative high-resolution snapshot of a reconstructed cluster of chromatin arrays in the 25 bp chromatin condensate. The top-scoring array (i.e. with greatest structural similarity to a traced array in a cryo-ET tomogram) is highlighted with DNA in dark pink and histones in light pink. All the arrays interacting with the top-scoring array are shown (17 arrays in total form the interaction cluster). One of the 16 arrays interacting with the top-scoring array is highlighted in orange. The inset expands the image of the two interacting arrays, to show face–to-face inter-array stacking. **(H)**As in panel G, but for a 30-bp chromatin condensates. The top-scoring array is highlighted with DNA in dark green and histones in light green, and the array interacting with it with DNA in dark cyan and histones in light cyan. All the arrays interacting with the top-scoring array are shown (12 arrays in total form the interaction cluster). The insets face-to–side inter-array interactions, and face-to–face intra-array contacts. **(I)**Number of inter-nucleosome contacts between histone tails of one nucleosome and the DNA (nucleosomal and linker) and histones (core and tails) of a neighboring nucleosome, quantified from molecular dynamics simulations of various different (Methods) reconstructed interacting clusters per condition (25 bp is magenta, 30 bp is green). Top panel shows intra-array contacts. Bottom panel shows inter-array contacts. Data for 25 bp fibers is shown in pink and for 30 bp fibers in green. The blue and cyan vertical lines show the positions of lysines and arginines, respectively. The shading indicates one standard deviation from the mean.

To understand how pairwise packing of nucleosomes generates interactions between arrays we next sought to connect the assigned nucleosomes into molecular units. Ambiguities in nucleosome positions/orientations and weak linker DNA density prevented us from connecting all nucleosomes into arrays to produce complete molecular representations of the condensates. Nevertheless, we were able to connect 10 groups of >8 nucleosomes into molecular units in the 25 bp and 30 bp condensates. For each molecule we also identified all surrounding nucleosomes (Fig. 5C, D); in most cases these neighbors could not be joined into molecular units themselves.

As illustrated in Figures 5C and S3D, within condensates the 25 bp nucleosome arrays are relatively expanded, with an average radius of gyration, Rg, of 18.2 ±1.3 nm. The molecules adopt numerous, irregular conformations. In contrast, the 30 bp arrays are compact, with average radius of gyration of 13.9 ± 0.9 nm, and composed mostly of groups of 4-6 nucleosomes stacked in zig-zag fashion (i.e., with nucleosomes N and N+2 interacting face-to-face) organized into two-start helices (Fig. 5D, S3D-F). For both chromatin types, the structures observed within the condensates have similar values and distributions of Rg, α, D and para to those observed in the dilute phase under the same solution conditions (fig. S3C, D, G-I), suggesting that phase separation may not strongly impact the conformation of individual molecules. One caveat to this interpretation is that we do not know whether molecules recently dissociated from the condensate and captured by freezing proximal to the liquid-liquid interface have relaxed to their equilibrium dilute-phase configurations.

The differences in molecular conformation produce starkly different patterns of intra- and intermolecular interactions for the two chromatin types (Fig. 5C, D). Due to their extended nature and the arrangements of successive nucleosomes, the 25 bp arrays show very few intramolecular nucleosome-nucleosome contacts and a large number of intermolecular contacts (Fig. 5C). The behavior is inverted for the 30 bp arrays, which make ∼4-fold more intramolecular contacts and ∼half as many intermolecular contacts than the 25 bp arrays. The types of intermolecular interactions made by the two systems are also different (Fig. 5E). Intermolecular contacts of the 25 bp arrays are dominated by energetically favorable (84, 85) face-to-face and face-to-side nucleosome geometries (Fig 5F). In contrast, the 30 bp arrays show fewer face-to-face and face-to-side geometries, and a greater proportion of energetically weaker (84, 85) side-to-side contacts (Fig 5F). Together these analyses reveal that within condensates 25 bp chromatin molecules make a larger number of intermolecular contacts, with geometries that are more energetically favorable, compared to 30 bp chromatin molecules. These differences suggest a greater enthalpic gain upon condensate formation for the 25 bp chromatin.

To understand how the histone tails contribute to intermolecular interactions between chromatin molecules within the condensates, we again turned to molecular simulations. As detailed in Methods and Figure S4A, starting with coarse grained simulations of condensates, we identified individual arrays whose conformations were nearly identical to molecules in the cryo-ET tomograms, and for each we added all the arrays in contact with it to produce a cluster. We then built chemically specific models of the clusters and used these to perform molecular dynamics simulations, with restraints applied to preserve the positions of all nucleosomes (Fig. 5G-H). Paralleling the intermolecular array-nucleosome contacts observed in the cryo-ET tomograms (Fig. 5E), the simulations revealed that in the 25-bp condensates, histone tails more frequently make inter-array interactions than intra-array interactions (Fig. 5I). In contrast, and again in agreement with the cryo-ET data (Fig. 5E), the majority of the histone tail interactions in the 30-bp condensates are intra-array (Fig. 5I).

In the 25-bp condensates, chromatin fibers interact frequently via their H4 and H2A(N) tails (Fig. 5I), consistent with the prevalence of inter-array face-to-face stacking (Fig. 3A, 5F-G). Conversely, within the 30-bp condensates, the H4 and H2A(N) tails play a more modest role in inter-array contacts (Fig. 5I), as they are primarily involved in bridging faceto-face contacts within their own fibers (Fig. 5H). This results in a relatively flat distribution of inter-array contacts across the four tails for the 30-bp chromatin. Notably, the pattern of inter-array contacts in 25 bp chromatin resembles that of intra-array contacts in 30 bp chromatin (dominant H4 and H2A(N)), since both are enriched in nucleosome faceto-face stacking. However, H2B interactions are numerous in the former, but are very few in the latter. This difference arises because antiparallel stacking in 25 bp chromatin positions the H2B tail in contact with the entry and exit DNA of the neighboring nucleosome, whereas parallel stacking in 30 bp chromatin positions it more peripherally. Thus, analogous to single fibers (Fig. 3), intermolecular contacts of the histone tails in condensates are determined by a combination of fiber conformation and tail position within the nucleosome. As described in Supplementary Tables 3-4, the patterns of tail interactions observed here are generally consistent with previous biochemical studies of tail contributions to chromatin oligomerization, with the caveat that for 30 bp chromatin, perturbations disrupting nucleosome stacking will affect both oligomerization and the conformation of individual arrays.

These data lead to a model in which linker length determines the conformation of chromatin molecules, which in turn determines the balance between intra- and intermolecular contacts involving nucleosome cores and histone tails as well as the energetic strength of those contacts. These contacts then define the thermodynamic stability of their condensates. Twenty-five bp chromatin adopts conformations favoring strong and numerous intermolecular interactions, producing more stable condensates. Thirty bp chromatin adopts conformations that allow weaker and fewer intermolecular interactions (but more and stronger intramolecular interactions), producing less stable condensates (Movie S1).

### Linker length determines connectivity of the condensate network

We next used the minimal model coarse-grained simulations to understand how the network of intermolecular interactions between chromatin arrays produces material properties of the condensates. We abstracted the molecules from simulations at selected timepoints onto a graph network where nodes represent individual nucleosome arrays and edges represent interactions between them (Fig. 6A, B, S5A, B). Inter-nucleosome contacts were weighted by the nucleosome pair orientations according to the potential of mean force (84). Figure 6A, B shows cross-sections through the centers of representative networks generated by simulations of 25 bp (left) and 30 bp (right) chromatin. The 25 bp network is more densely connected than the 30 bp network, as described quantitatively by the average number (fig. S5C) and strength (fig. S5D and E) of pairwise inter-array interactions, and normalized algebraic connectivity values of 0.67 ± 0.03 and 0.53 ± 0.03 (on a scale of 0 – 1, where 0 indicates no connections and 1 indicates all nodes connected with the highest node degree in the 25 bp chromatin), respectively.

**Figure 6.**
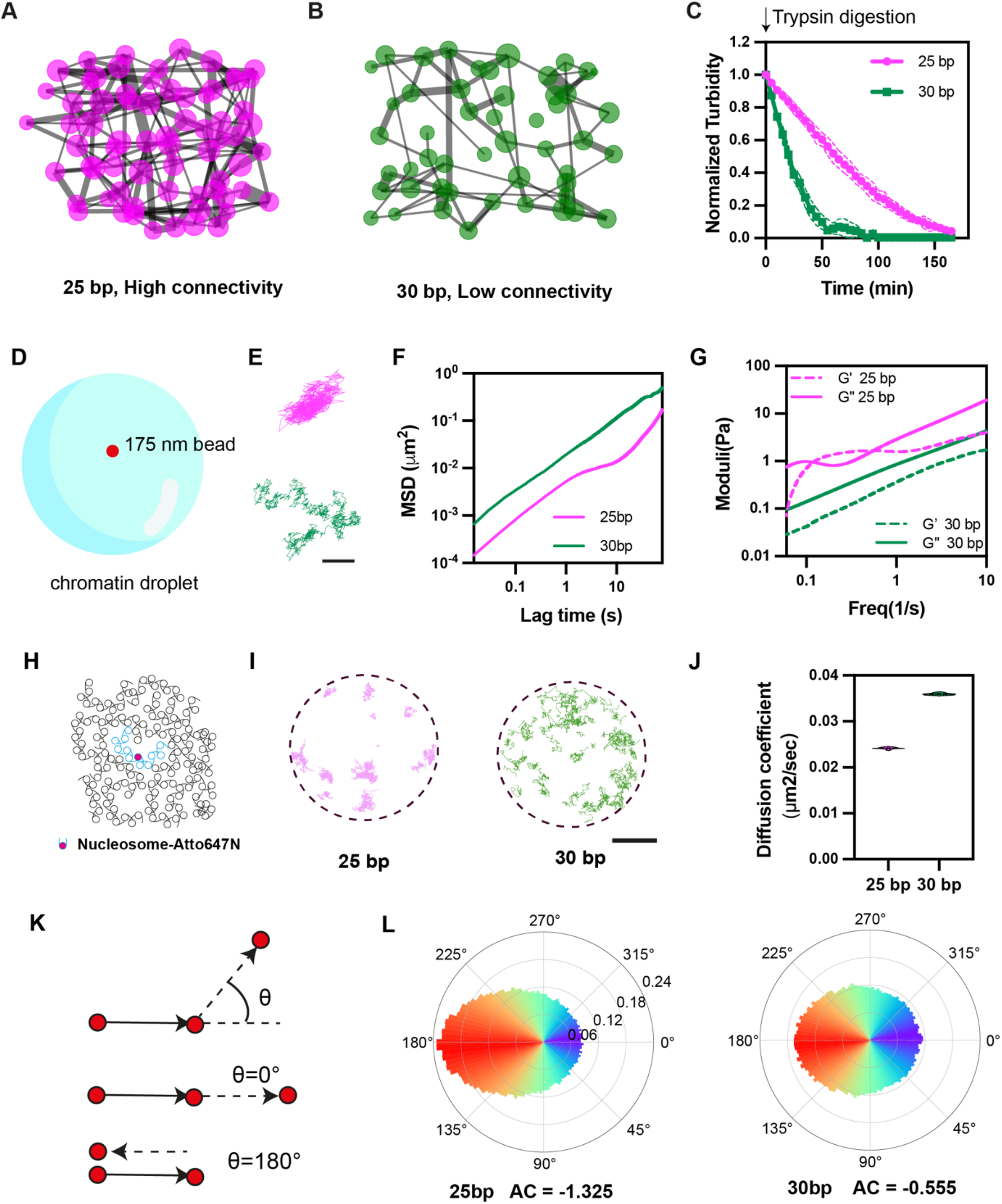
Distinct dynamics of 25 bp and 30 bp chromatin condensate on molecularand meso-scales. **(A-B)** Cross sections of interaction networks derived from coarse-grained simulations of 25 bp (A) and 30 bp (B) chromatin condensates. Node diameter is scaled by the number of molecules contacted by each array and edge thickness is scaled by the energy of association between each pair of molecules. **(B)**Normalized turbidity (A345 nm) of solutions containing chromatin droplets after introduction of trypsin, which cleaves histone tails. **(C)**Schematic illustration of particle tracking microrheology. Fluorescent 175 nm beads are introduced into condensates, and their random movement is tracked over time using fluorescence microscopy. **(D)**Representative trajectories of particles tracked within 25 bp (top) and 30 bp (bottom) chromatin condensates. Scale bar is 100 nm. **(E)**Mean Squared Displacement (MSD) analysis for beads tracked in 25 bp (magenta) and 30 bp (green) chromatin condensates. **(F)**Elastic/storage (G’) and viscous/loss (G’’) moduli of 25 bp (magenta) and 30 bp (green) chromatin condensates, calculated from MSD data in panel (F). **(G)**Schematic illustration of single-molecule tracking experiments. Sparse labelling of nucleosome arrays with Atto647N dye enables precise visualization of the dynamics of individual molecules. **(H)**Trajectories of single nucleosome array molecules in 25 bp (magenta) and 30 bp (green) chromatin condensates. Scale bar = 1 μm. **(I)**Diffusion coefficients calculated for nucleosome arrays in 25 bp and 30 bp chromatin condensates. **(J)**Schematic illustration of trajectory angular analysis, where the angle between every pair of steps in a single-molecule trajectory is measured. Zero degrees corresponds to directed motion and 180 degrees represents back-and-forth movement. **(K)**Trajectory angular analysis distributions for nucleosome arrays in 25 bp (left) and 30 bp (right) chromatin condensates. Asymmetry coefficient (AC) describes the ratio of forward to backward movement on a log2 scale

To examine network connectivity biochemically we treated the condensates with trypsin, which cleaves histone tails leading to dissolution ((33), (Fig. 6C)). While 30 bp condensates dissolved quickly upon treatment with trypsin, 25 bp condensates were more resistant, consistent with greater histone tail-mediated connectivity. We note, however, that we cannot rule out additional effects due to changes in molecular structure, which might secondarily affect phase separation, due to removal of the histone tails. We also modeled this process by reducing the number of potential interactions between nucleosomes in coarse grained simulations. We found that the size of the largest connected cluster of molecules decreased much faster for 30 bp chromatin compared to 25 bp chromatin, aligning with the experimental results (fig. S5F).

Thus, the differences in molecular interactions of 25 bp and 30 bp chromatin produce higher order networks with different degrees of connectivity within condensates (Movie S1).

### 25 bp and 30 bp chromatin show distinct dynamics on the meso- and molecular scales

The differences in affinity (more precisely, lifetime) and density of bonds in 25 bp and 30 bp chromatin condensates suggested that the material properties of the two liquids might differ (5, 30, 107, 108). To test this idea, we initially measured mesoscale dynamics of the condensates through particle tracking microrheology, which reports on macroscopic viscoelasticity (Fig. 6D). Figure 6E shows representative trajectories of 175 nm-diameter fluorescent beads undergoing Brownian motion within 25 bp (top) and 30 bp (bottom) chromatin condensates, respectively. The beads sample smaller volumes in the former system, revealing more constrained motions. Analysis of mean-squared displacement (MSD) versus lag time revealed that beads show purely diffusive (viscous) behavior (linear MSD vs lag time with slope ∼1) in 30 bp chromatin condensates over timescales of ∼0.015 – 150 seconds (Fig. 6F). In contrast, the 25 bp chromatin droplets showed lower mean-squared displacement values on these timescales, and strong non-linearity in the ∼1-10 second regime, indicating sub-diffusive motion and viscoelasticity. We modeled the condensates as Maxwell fluids and used the generalized Stokes-Einstein equation to calculate their viscoelastic moduli (G*), composed of viscous (G”) and elastic (G’) components. As suggested by the mean-squared displacement behavior, 25 bp chromatin condensates exhibited higher viscous and elastic components than 30 bp chromatin condensates in all measured time regimes (Fig. 6E, F). Moreover, while for the 30 bp condensates viscosity is dominant (G” > G’) on all timescales (Fig. 6G), for the 25 bp condensates the viscous and elastic components intersect at 0.12 Hz and again at 0.56 Hz (Fig. 6G). Within this frequency regime, elasticity is dominant (G’ > G”). Thus, the greater connectivity and likely longer-lived intermolecular interactions of the 25 bp chromatin produce greater overall viscoelasticity (G*), and qualitatively different relationships between viscosity and elasticity (G” vs G’), than 30 bp chromatin. These results show how viscoelastic behaviors are determined by the molecular architecture of chromatin condensates, which in turn affect the meso-scale dynamics of large internal components.

We also examined dynamics of the condensates on molecular scales using single-molecule imaging. We sparsely labelled histones (∼1:10,000) with Atto 647N and tracked the movement of single arrays within condensates using highly inclined and laminated optical sheet (HILO) microscopy (Fig. 6H). Single 25 bp arrays exhibited slower dynamics than 30 bp arrays, with diffusion coefficients of 0.024±0.0002 µm2/sec and 0.036 ±0.0002 µm2/sec, respectively (Fig. 6I and 6J). This result aligns with the more numerous and stronger interactions of 25 bp arrays than 30 bp arrays within their respective condensates (Fig. 5E). Additionally, the distribution of jump angles between two consecutive translocations in the trajectories (Fig. 6K) was significantly biased toward 180° for 25 bp arrays compared to 30 bp arrays, indicating less random and more confined dynamics (Fig. 6L). Notably, the calculated asymmetry coefficient (AC), which describes the ratio of forward to backward movement on a log2 scale, of 25 bp arrays is similar to that previously reported for single nucleosome dynamics in live cells (−1.33 versus -1.36 (109)). These data suggest that the connectivity and interaction dynamics of the 25 bp condensates constrain the motion of individual molecules relative to that observed in 30 bp chromatin.

Together, the rheological and single-molecule data suggest that differences in network connectivity and interaction affinity (lifetime), which arise from differences in linker-dependent molecular structure, produce quantitative and qualitative differences in the dynamics of chromatin condensates spanning molecular to meso-scales (Movie S1).

### Native chromatin forms discrete domains with nucleosome organization similar to 25 bp condensates

We investigated chromatin organization in vitrified HeLa cell nuclei and intact NIH 3T3 cells using cryo-FIB milling and cryo-ET (Fig. 7A, B). Each system had nuclear domains of high nucleosome density, varying in size from ∼100-300 nm, surrounded by regions of lower nucleosome density. Some of these domains were clearly discrete (Fig. 7C, F). Others appeared discrete in two-dimensional sections but were connected when analyzed in three dimensions. These structures, which were also reported in a recent cryoET study of T-lymphoblast CEM RPE-1 nuclei (65, 70), are similar in size to domains of high-density DNA observed by super-resolution imaging (49, 50). Staining for epigenetic marks suggested that those structures corresponded to inactive chromatin (50). Thus, it is possible that the high-density domains we observe here are inactive chromatin as well.

**Figure 7.**
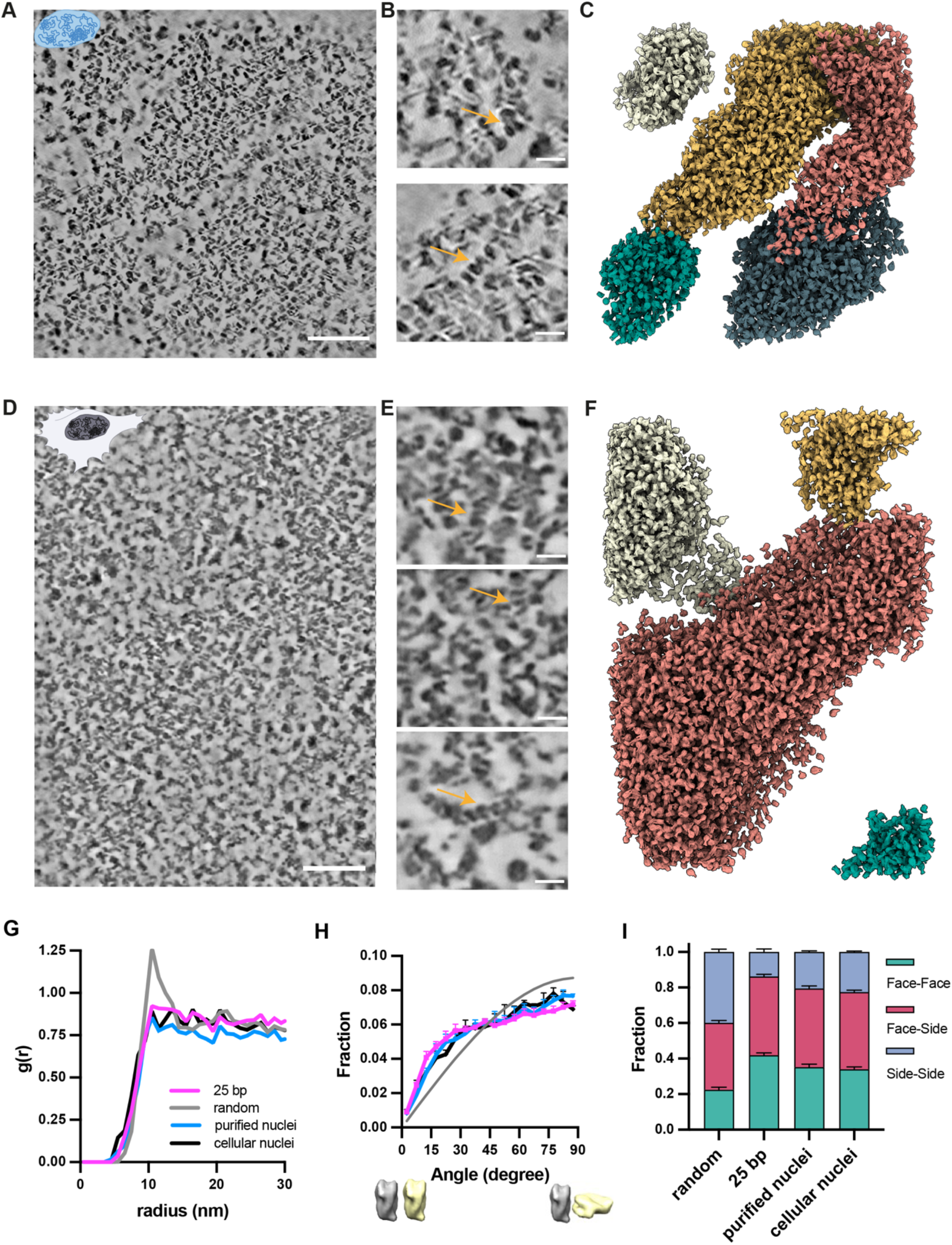
Native chromatin forms discrete domains and is organized similarly to 25 bp chromatin condensates. **(A-F)** Tomographic slices and 3D reconstructions illustrating nuclear nucleosome organization. (A-C) Overview slice (scale bar = 100 nm), expanded regions (scale bar = 20 nm) and 3D nucleosome model assignments from a purified Hela cell nucleus. (D-F) Analogous images from the nucleus of an intact NIH3T3 cell. Note that panels B and E are not expansions of regions from A and D, but are derived from different planes of the tomograms. **(G)**Radial distribution functions, g(r), for randomly oriented nucleosomes (grey), nucleosomes in 25 bp chromatin condensates (magenta), purified Hela cell nuclei (cyan), and nuclei of intact NIH3T3 cells (black). **(H)**Distributions of nearest di-nucleosome orientations for 25 bp chromatin condensates (magenta), purified Hela cell nuclei (cyan), and nuclei of intact NIH3T3 cells (black). Zero and 90 degrees represent parallel and perpendicular orientations, respectively. **(I)**Fractions of face-to-face, face-to-side and side-to-side pairwise nucleosome contacts in a random nucleosome distribution, 25 bp reconstituted chromatin condensates, nucleosomes in purified Hela cell nuclei, and nuclei of intact NIH3T3 cells.

Radial distribution functions of ∼20,000 nucleosomes in each native chromatin sample were similar to those of the synthetic 25 bp chromatin condensates, rising from ∼6 nm to a weak maximum at ∼11 nm and a broad plateau at longer distances (Fig. 7G). The pairwise orientation distribution of nearest nucleosome pairs was also similar for native chromatin and the 25 bp chromatin condensates, with all showing small deviations from a random distribution (Fig. 7H). The native chromatin and 25 bp chromatin condensates also showed a similar distribution of nucleosome packing geometries, which differed from a random bath of nucleosomes by an excess of face-to-face/side and depletion of side-to-side contacts (Fig. 7I). Thus, at the level of individual nucleosomes, dense chromatin regions and 25 bp chromatin condensates have packing arrangements that deviate similarly from randomly organized molecules, likely reflecting geometric features of nucleosome interaction energies and restrictions imparted by rigid DNA linkers.

Inspection of the dense chromatin regions revealed some groups of stacked nucleosomes (Fig. 7B, E). These form two-start helices analogous to those in the synthetic 30 bp chromatin condensates, suggesting they may be formed by genomic regions with nucleosomes phased with ∼10N spacing. These structures could be functionally important if recognized by specific factors or associated with specific loci. Overall, our data show that native chromatin from these cellular sources forms ∼100-300 nm scale dense domains composed mostly of nucleosomes that are packed non-randomly and have an excess of contacts involving nucleosome faces, with occasional instances of two-start helical fibers.

## Discussion

Our studies of chromatin illustrate the types of information that may be gained from future cryo-ET studies of other biomolecular condensates. In cases where condensate formation is induced by changes in environmental conditions, comparison of dilute phase structures above and below the phase separation threshold can provide insight into the conformational changes that drive assembly. Analysis of internal molecules can reveal the hierarchy of structures and interactions that compose and dictate the properties of the condensate. Sub-tomogram averaging can afford medium resolution (∼6 Å in the present systems) structures of individual particles. For multivalent systems, depending on the degree of molecular flexibility, it may be possible to reconstruct larger order units of the core particle, analogous to the di-, tri- and tetra-nucleosome groupings we could visualize here for 30 bp chromatin. It may be possible to observe and reconstruct client molecules (which bind the phase separating scaffold molecules) in cryo-ET tomograms. Such studies could reveal how clients are affected by a condensate, bind to scaffolds, and modulate condensate architecture. At a higher level, one can visualize the spatial arrangement of components on nano-to micrometer length scales. Such analyses will reveal local packing interactions (distances, geometries; energetics, when coupled with modeling), as well as the networks that arise from these contacts. Network structure will in turn reveal the distribution of pore sizes. When coupled with dynamics measurements, the network structure will also inform on viscoelasticity. While not explored here, comparison of molecular organization in the core and surface of a condensate could provide a structural explanation for surface tension and potentially surface chemistry.

These features are relevant to diverse aspects of condensate function. Depending on resolution, structures of individual particles within a condensate may provide insight into changes in activity that accompany phase separation. Pore sizes in the molecular network of a condensate will be relevant to recruitment and movement of other molecules within the compartment, again with relevance to internal chemistry (110–113). Relatedly, viscoelasticity will inform on the length and timescales at which internal processes will be impacted, and how the condensate will respond to internal and external mechanical forces (114–116). When coupled with simulations, the organization of macromolecules within a condensate, and its time-dependent changes, will provide a structural framework to understand the physical properties of the compartment such as pH, ion concentrations (15, 117) and hydrophobicity (14, 118). Finally, condensate structural analyses should explain how disease-causing mutations alter both molecular and higher order organization to produce functional defects (119). Importantly, it remains unknown which aspects of condensate function derive from the structure and activity of individual building blocks and which derive from higher-order assembly. As in the understanding of macromolecular machines through canonical structural analyses, further studies of condensate structure coupled with specific perturbations and quantitative functional read outs will be necessary to reveal structure-function relationships in these micrometer scale systems.

Understanding the structural and functional heterogeneity of cellular chromatin is important in understanding genome function. Our cryo-ET images here, and from others (65, 70), show that nuclear chromatin forms dense ∼100-300 nanometer foci of concentrated nucleosomes with relatively clear boundaries (especially in the highest contrast images, Fig. 7A), surrounded by regions of appreciably lower density. Both these foci and our 25 bp chromatin condensates have pairwise nucleosome arrangements and packing geometries that are similar, but not identical to those expected for a random distribution. It is possible that the foci form through micro-phase separation (i.e. phase separation of a small element of one or more chromosomes (120)), although other mechanisms are also possible. One consequence of a phase separation model is that chromatin could be switched sharply between states of density and function by changes in inter-nucleosome interaction strength. Our findings here, as well as recent biochemical data (84) suggest this could be achieved by nucleosome remodelers, which, by moving nucleosomes along DNA filaments, could control both local chromatin structure as well as higher-order organization and dynamics. It could also be achieved by factors that affect nucleosome contacts, including posttranslational modifications such as acetylation or ubiquitination that alter binding interactions of histone tails (121). Together, these factors could contribute to the heterogeneity of chromatin that exists in cells.

It remains unclear how the foci observed by cryo-ET relate to genome organization characterized by single-molecule imaging and interaction mapping technologies. The former have shown that small groups of 10-50 nucleosomes, termed clutches, assemble into discrete units (36). The latter have suggested that on a cell population basis genomic regions of 0.2 – 1 Mbp associate into Topologically Associated Domains (TADs), which show numerous self-interactions but are largely insulated from surrounding regions (36, 122, 123). Such large domains are not typically observed in interaction maps of individual cells sampled at single time points, where only subsets appear to associate (124–127). But the larger TADs may reflect cellular genome organization averaged over time. The foci we observe by cryo-ET have density of approximately 600,000 nucleosomes/µm3. Assuming an average nucleosome repeat length of 192 base pairs (128), this density would correspond to 115 Mb base pairs/µm3. As a rough estimate, foci of 100-300 nm diameter would thus contain 315 – 8500 nucleosomes and 60 kbp – 1.6 Mbp of DNA. Thus, the foci are much larger than clutches and more consistent in size with TADs. It remains unknown how these foci, each of which represents only a tiny fraction of the human genome (< 0.5 %), relate to nuclear functions such as transcription, replication, etc. Future studies based on correlated lightand electron microscopy, to understand epigenetic marks and perhaps genomic regions present in the foci, will be important to understanding how these higher-order chromatin structures form and what their functions may be.

## Acknowledgments

We thank Daniel and David Isenberg for their generous support of the MBL Chromatin Collaborative in honor of their father, Irvin Isenberg, and the other members of the Collaborative for critical insights and discussion. We thank Sy Redding and Christopher Woodilla for assistance with and discussions regarding chromatin viscoelasticity experiments. We thank Janet Iwasa for support and advice during early stages of visualizing chromatin condensates.

## Funding

Howard Hughes Medical Institute (M.K.R., Z.Y., and E.V.); Paul G. Allen Frontiers Distinguished Investigator Award (M.K.R.); Welch Foundation I-1544 (M.K.R.); European Research Council (ERC) under the European Union’s Horizon 2020 research and innovation program, grant agreement no. 803326 (R.C.G). Cancer Research and Prevention Institute of Texas RP220582 (UTSW cryo-EM Facility, Structure Biology Lab); NIH Common Fund Transformative High Resolution Cryo-Electron Microscopy program U24 GM129539 (National Center For In-Situ Tomographic Ultramicroscopy (NCITU)); Engineering & Physical Sciences Research Council (EPSRC) EP/R029407/1 (R.C.G.)

## Author contributions

Conceptualization: HZ, MKR

Methodology: HZ, JH, MJM, KR,JHH, RY, RCG, MKR

Investigation: HZ, JH, MJM, KR, JHH, RY, JH, MS, XZ, LKD, BAG, JRE

Visualization: HZ, JH, MR

Funding acquisition: EV, RCG, MKR

Project administration: ZH, EV, RCG, MKR

Supervision: EV, RCG, MKR

Writing – original draft: HZ, MKR

Writing – review & editing: all authors

## Competing interests

Authors declare that they have no competing interests.

## Data and materials availability

All data generated during the current study are available from the corresponding authors on request from the referees, and will be posted to the Dryad database before publication.

The CATM software, and all analysis scripts will be available on the Rosen lab Gitlab site before publication, and have been provided to referees with the manuscript. All simulation code will be deposited in Collepardo lab Gitlab.

All reagents described in the work will be available by request to the corresponding authors.

## Supplementary Materials

Materials and Methods

Supplementary Text

Figs. S1 to S5

Tables S1 to S5

Movies S1

References

## Notes

### Competing Interest Statement

The authors have declared no competing interest.

## References

1. P. Li, S. Banjade, H.-C. Cheng, S. Kim, B. Chen, L. Guo, M. Llaguno, J. Hollingsworth, D. S. King, S. F. Banani, P. S. Russo, Q.-X. Jiang, B. T. Nixon, M. K. Rosen, Phase transitions in the assembly of multivalent signalling proteins. Nature 483, 336–340 (2012).

2. S. F. Banani, H. O. Lee, A. A. Hyman, M. K. Rosen, Biomolecular condensates: organizers of cellular biochemistry. Nature Reviews Molecular Cell Biology 18, 285–298 (2017).

3. Y. Shin, C. P. Brangwynne, Liquid phase condensation in cell physiology and disease. Science 357, eaaf4382 (2017).

4. I. Alshareedah, W. M. Borcherds, S. R. Cohen, A. Singh, A. E. Posey, M. Farag, A. Bremer, G. W. Strout, D. T. Tomares, R. V. Pappu, T. Mittag, P. R. Banerjee, Sequence-specific interactions determine viscoelasticity and ageing dynamics of protein condensates. Nat. Phys., doi: 10.1038/s41567-024-02558-1 (2024).

5. S. R. Cohen, P. R. Banerjee, R. V. Pappu, Direct computations of viscoelastic moduli of biomolecular condensates. The Journal of Chemical Physics 161, 095103 (2024).

6. A. Chandrasekaran, K. Graham, J. C. Stachowiak, P. Rangamani, Kinetic trapping organizes actin filaments within liquid-like protein droplets. Nat Commun 15, 3139 (2024).

7. A. S. Lyon, W. B. Peeples, M. K. Rosen, A framework for understanding the functions of biomolecular condensates across scales. Nat Rev Mol Cell Biol 22, 215–235 (2021).

8. D. L. J. Lafontaine, J. A. Riback, R. Bascetin, C. P. Brangwynne, The nucleolus as a multiphase liquid condensate. Nat Rev Mol Cell Biol 22, 165–182 (2021).

9. A. Yamasaki, J. Md. Alam, D. Noshiro, E. Hirata, Y. Fujioka, K. Suzuki, Y. Ohsumi, N. N. Noda, Liquidity Is a Critical Determinant for Selective Autophagy of Protein Condensates. Molecular Cell 77, 1163–1175.e9 (2020).

10. Z. Wang, D. Chen, D. Guan, X. Liang, J. Xue, H. Zhao, G. Song, J. Lou, Y. He, H. Zhang, Material properties of phase-separated TFEB condensates regulate the autophagy-lysosome pathway. Journal of Cell Biology 221, e202112024 (2022).

11. P. Guo, B. Li, W. Dong, H. Zhou, L. Wang, T. Su, C. Carl, Y. Zheng, Y. Hong, H. Deng, D. Pan, PI4P-mediated solid-like Merlin condensates orchestrate Hippo pathway regulation. Science 385, eadf4478 (2024).

12. J. Risso-Ballester, M. Galloux, J. Cao, R. Le Goffic, F. Hontonnou, A. Jobart-Malfait, A. Desquesnes, S. M. Sake, S. Haid, M. Du, X. Zhang, H. Zhang, Z. Wang, V. Rincheval, Y. Zhang, T. Pietschmann, J.-F. Eléouët, M.-A. Rameix-Welti, R. Altmeyer, A condensate-hardening drug blocks RSV replication in vivo. Nature 595, 596–599 (2021).

13. L.-P. Bergeron-Sandoval, S. Kumar, H. K. Heris, C. L. A. Chang, C. E. Cornell, S. L. Keller, P. François, A. G. Hendricks, A. J. Ehrlicher, R. Pappu, S. W. Michnick, Endocytic proteins with prion-like domains form viscoelastic condensates that enable membrane remodeling. Proc. Natl. Acad. Sci. U.S.A. 118, e2113789118 (2021).

14. S. Ambadi Thody, H. D. Clements, H. Baniasadi, A. S. Lyon, M. S. Sigman, M. K. Rosen, Small-molecule properties define partitioning into biomolecular condensates. Nat. Chem., doi: 10.1038/s41557-024-01630-w (2024).

15. M. R. King, K. M. Ruff, A. Z. Lin, A. Pant, M. Farag, J. M. Lalmansingh, T. Wu, M. J. Fossat, W. Ouyang, M. D. Lew, E. Lundberg, M. D. Vahey, R. V. Pappu, Macromolecular condensation organizes nucleolar sub-phases to set up a pH gradient. Cell 187, 1889–1906.e24 (2024).

16. A. E. Posey, A. Bremer, N. A. Erkamp, A. Pant, T. P. J. Knowles, Y. Dai, T. Mittag, R. V. Pappu, Biomolecular Condensates are Characterized by Interphase Electric Potentials. J. Am. Chem. Soc., jacs.4c08946 (2024).

17. E. W. Martin, C. Iserman, B. Olety, D. M. Mitrea, I. A. Klein, Biomolecular Condensates as Novel Antiviral Targets. Journal of Molecular Biology 436, 168380 (2024).

18. D. M. Mitrea, M. Mittasch, B. F. Gomes, I. A. Klein, M. A. Murcko, Modulating biomolecular condensates: a novel approach to drug discovery. Nat Rev Drug Discov 21, 841–862 (2022).

19. T. H. Kim, B. Tsang, R. M. Vernon, N. Sonenberg, L. E. Kay, J. D. Forman-Kay, Phospho-dependent phase separation of FMRP and CAPRIN1 recapitulates regulation of translation and deadenylation. Science 365, 825–829 (2019).

20. E. W. Martin, A. S. Holehouse, I. Peran, M. Farag, J. J. Incicco, A. Bremer, C. R. Grace, A. Soranno, R. V. Pappu, T. Mittag, Valence and patterning of aromatic residues determine the phase behavior of prion-like domains.

21. A. C. Murthy, G. L. Dignon, Y. Kan, G. H. Zerze, S. H. Parekh, J. Mittal, N. L. Fawzi, Molecular interactions underlying liquid−liquid phase separation of the FUS low-complexity domain. Nat Struct Mol Biol 26, 637–648 (2019).

22. M. Bose, M. Lampe, J. Mahamid, A. Ephrussi, Liquid-to-solid phase transition of oskar ribonucleoprotein granules is essential for their function in Drosophila embryonic development. Cell 185, 1308–1324.e23 (2022).

23. M. Zhang, C. Díaz-Celis, B. Onoa, C. Cañari-Chumpitaz, K. I. Requejo, J. Liu, M. Vien, E. Nogales, G. Ren, C. Bustamante, Molecular organization of the early stages of nucleosome phase separation visualized by cryo-electron tomography. Molecular Cell, S1097276522006505 (2022).

24. J. Guillén-Boixet, A. Kopach, A. S. Holehouse, S. Wittmann, M. Jahnel, R. Schlüßler, K. Kim, I. R. E. A. Trussina, J. Wang, D. Mateju, I. Poser, S. Maharana, M. Ruer-Gruß, D. Richter, X. Zhang, Y.-T. Chang, J. Guck, A. Honigmann, J. Mahamid, A. A. Hyman, R. V. Pappu, S. Alberti, T. M. Franzmann, RNA-Induced Conformational Switching and Clustering of G3BP Drive Stress Granule Assembly by Condensation. Cell 181, 346–361.e17 (2020).

25. F. Tollervey, X. Zhang, M. Bose, J. Sachweh, J. B. Woodruff, T. M. Franzmann, J. Mahamid, “Cryo-Electron Tomography of Reconstituted Biomolecular Condensates” in Phase-Separated Biomolecular Condensates: Methods and Protocols, H.-X. Zhou, J.-H. Spille, P. R. Banerjee, Eds. (Springer US, New York, NY, 2023; 10.1007/978-1-0716-2663-4_15), pp. 297–324.

26. X. Zhang, S. Sridharan, I. Zagoriy, C. Eugster Oegema, C. Ching, T. Pflaesterer, H. K. H. Fung, I. Becher, I. Poser, C. W. Müller, A. A. Hyman, M. M. Savitski, J. Mahamid, Molecular mechanisms of stress-induced reactivation in mumps virus condensates. Cell 186, 1877–1894.e27 (2023).

27. J. B. Woodruff, B. Ferreira Gomes, P. O. Widlund, J. Mahamid, A. Honigmann, A. A. Hyman, The Centrosome Is a Selective Condensate that Nucleates Microtubules by Concentrating Tubulin. Cell 169, 1066–1077.e10 (2017).

28. F. J. B. Bäuerlein, I. Saha, A. Mishra, M. Kalemanov, A. Martínez-Sánchez, R. Klein, I. Dudanova, M. S. Hipp, F. U. Hartl, W. Baumeister, R. Fernández-Busnadiego, In Situ Architecture and Cellular Interactions of PolyQ Inclusions. Cell 171, 179–187.e10 (2017).

29. X. Liu, X. Xia, M. W. Martynowycz, T. Gonen, Z. H. Zhou, Molecular sociology of virus-induced cellular condensates supporting reovirus assembly and replication. Nat Commun 15, 10638 (2024).

30. A. R. Tejedor, R. Collepardo-Guevara, J. Ramírez, J. R. Espinosa, Time-Dependent Material Properties of Aging Biomolecular Condensates from Different Viscoelasticity Measurements in Molecular Dynamics Simulations. J. Phys. Chem. B 127, 4441–4459 (2023).

31. A. R. Tejedor, I. Sanchez-Burgos, M. Estevez-Espinosa, A. Garaizar, R. Collepardo-Guevara, J. Ramirez, J. R. Espinosa, Protein structural transitions critically transform the network connectivity and viscoelasticity of RNA-binding protein condensates but RNA can prevent it. Nat Commun 13, 5717 (2022).

32. I. Sanchez-Burgos, J. A. Joseph, R. Collepardo-Guevara, J. R. Espinosa, Size conservation emerges spontaneously in biomolecular condensates formed by scaffolds and surfactant clients. Sci Rep 11, 15241 (2021).

33. B. A. Gibson, L. K. Doolittle, M. W. G. Schneider, L. E. Jensen, N. Gamarra, L. Henry, D. W. Gerlich, S. Redding, M. K. Rosen, Organization of Chromatin by Intrinsic and Regulated Phase Separation. Cell 179, 470–484.e21 (2019).

34. M. W. G. Schneider, B. A. Gibson, S. Otsuka, M. F. D. Spicer, M. Petrovic, C. Blaukopf, C. C. H. Langer, P. Batty, T. Nagaraju, L. K. Doolittle, M. K. Rosen, D. W. Gerlich, A mitotic chromatin phase transition prevents perforation by microtubules. Nature, doi: 10.1038/s41586-022-05027-y (2022).

35. B. A. Gibson, C. Blaukopf, T. Lou, L. Chen, L. K. Doolittle, I. Finkelstein, G. J. Narlikar, D. W. Gerlich, M. K. Rosen, In diverse conditions, intrinsic chromatin condensates have liquid-like material properties. Proc. Natl. Acad. Sci. U.S.A. 120, e2218085120 (2023).

36. M. Lakadamyali, M. P. Cosma, Visualizing the genome in high resolution challenges our textbook understanding. Nat Methods 17, 371–379 (2020).

37. A. G. Larson, D. Elnatan, M. M. Keenen, M. J. Trnka, J. B. Johnston, A. L. Burlingame, D. A. Agard, S. Redding, G. J. Narlikar, Liquid droplet formation by HP1α suggests a role for phase separation in heterochromatin. Nature 547, 236–240 (2017).

38. A. R. Strom, A. V. Emelyanov, M. Mir, D. V. Fyodorov, X. Darzacq, G. H. Karpen, Phase separation drives heterochromatin domain formation. Nature 547, 241–245 (2017).

39. I. Solovei, M. Kreysing, C. Lanctôt, S. Kösem, L. Peichl, T. Cremer, J. Guck, B. Joffe, Nuclear Architecture of Rod Photoreceptor Cells Adapts to Vision in Mammalian Evolution. Cell 137, 356–368 (2009).

40. H. Belaghzal, T. Borrman, A. D. Stephens, D. L. Lafontaine, S. V. Venev, Z. Weng, J. F. Marko, J. Dekker, Liquid chromatin Hi-C characterizes compartment-dependent chromatin interaction dynamics. Nat Genet 53, 367–378 (2021).

41. J. Dekker, L. A. Mirny, The chromosome folding problem and how cells solve it. Cell 187, 6424–6450 (2024).

42. R. R. Cheng, V. G. Contessoto, E. Lieberman Aiden, P. G. Wolynes, M. Di Pierro, J. N. Onuchic, Exploring chromosomal structural heterogeneity across multiple cell lines. eLife 9, e60312 (2020).

43. Y. Itoh, E. J. Woods, K. Minami, K. Maeshima, R. Collepardo-Guevara, Liquid-like chromatin in the cell: What can we learn from imaging and computational modeling? Current Opinion in Structural Biology 71, 123–135 (2021).

44. K. Luger, A. W. Mäder, R. K. Richmond, D. F. Sargent, T. J. Richmond, Crystal structure of the nucleosome core particle at 2.8 Å resolution. Nature 389, 251–260 (1997).

45. S. Pepenella, K. J. Murphy, J. J. Hayes, Intra- and inter-nucleosome interactions of the core histone tail domains in higher-order chromatin structure. Chromosoma 123, 3–13 (2014).

46. M. A. Ricci, C. Manzo, M. F. García-Parajo, M. Lakadamyali, M. P. Cosma, Chromatin Fibers Are Formed by Heterogeneous Groups of Nucleosomes In Vivo. Cell 160, 1145–1158 (2015).

47. M. Ohno, T. Ando, D. G. Priest, V. Kumar, Y. Yoshida, Y. Taniguchi, Sub-nucleosomal Genome Structure Reveals Distinct Nucleosome Folding Motifs. Cell 176, 520–534.e25 (2019).

48. H. D. Ou, S. Phan, T. J. Deerinck, A. Thor, M. H. Ellisman, C. C. O’Shea, ChromEMT: Visualizing 3D chromatin structure and compaction in interphase and mitotic cells. Science 357, eaag0025 (2017).

49. T. Nozaki, R. Imai, M. Tanbo, R. Nagashima, S. Tamura, T. Tani, Y. Joti, M. Tomita, K. Hibino, M. T. Kanemaki, K. S. Wendt, Y. Okada, T. Nagai, K. Maeshima, Dynamic Organization of Chromatin Domains Revealed by Super-Resolution Live-Cell Imaging. Molecular Cell 67, 282–293.e7 (2017).

50. E. Miron, R. Oldenkamp, J. M. Brown, D. M. S. Pinto, C. S. Xu, A. R. Faria, H. A. Shaban, J. D. P. Rhodes, C. Innocent, S. de Ornellas, H. F. Hess, V. Buckle, L. Schermelleh, Chromatin arranges in chains of mesoscale domains with nanoscale functional topography independent of cohesin. Science Advances 6, eaba8811 (2020).

51. G. Pei, H. Lyons, P. Li, B. R. Sabari, Transcription regulation by bio-molecular condensates. Nat Rev Mol Cell Biol, doi: 10.1038/s41580-024-00789-x (2024).

52. S. Brahmachari, S. Tripathi, J. N. Onuchic, H. Levine, Nucleosomes play a dual role in regulating transcription dynamics. Proc. Natl. Acad. Sci. U.S.A. 121, e2319772121 (2024).

53. S. Sekine, H. Ehara, T. Kujirai, H. Kurumizaka, Structural perspectives on transcription in chromatin. Trends in Cell Biology, S0962892423001551 (2023).

54. A. E. Ehrenhofer-Murray, Chromatin dynamics at DNA replication, transcription and repair. European Journal of Biochemistry 271, 2335–2349 (2004).

55. E. F. Hammonds, M. C. Harwig, E. A. Paintsil, E. A. Tillison, R. B. Hill, E. A. Morrison, Histone H3 and H4 tails play an important role in nucleosome phase separation. Biophysical Chemistry 283, 106767 (2022).

56. X. Li, Z. An, W. Zhang, F. Li, Phase Separation: Direct and Indirect Driving Force for High-Order Chromatin Organization. Genes 14, 499 (2023).

57. S. Sanulli, M. J. Trnka, V. Dharmarajan, R. W. Tibble, B. D. Pascal, A. L. Burlingame, P. R. Griffin, J. D. Gross, G. J. Narlikar, HP1 reshapes nucleosome core to promote phase separation of heterochromatin. Nature 575, 390–394 (2019).

58. H. Strickfaden, T. O. Tolsma, A. Sharma, D. A. Underhill, J. C. Hansen, M. J. Hendzel, Condensed Chromatin Behaves like a Solid on the Mesoscale In Vitro and in Living Cells. Cell 183, 1772–1784.e13 (2020).

59. A. Shakya, S. Park, N. Rana, J. T. King, Liquid-Liquid Phase Separation of Histone Proteins in Cells: Role in Chromatin Organization. Biophysical Journal 118, 753–764 (2020).

60. Q. Chen, L. Zhao, A. Soman, A. Y. Arkhipova, J. Li, H. Li, Y. Chen, X. Shi, L. Nordenskiöld, Chromatin Liquid–Liquid Phase Separation (LLPS) Is Regulated by Ionic Conditions and Fiber Length. Cells 11, 3145 (2022).

61. T. Nozaki, S. Shinkai, S. Ide, K. Higashi, S. Tamura, M. A. Shimazoe, M. Nakagawa, Y. Suzuki, Y. Okada, M. Sasai, S. Onami, K. Kurokawa, S. Iida, K. Maeshima, Condensed but liquid-like domain organization of active chromatin regions in living human cells. Sci. Adv. 9, eadf1488 (2023).

62. Y. Li, H. Zhang, X. Li, W. Wu, P. Zhu, Cryo-ET study from in vitro to in vivo revealed a general folding mode of chromatin with two-start helical architecture. Cell Reports 42, 113134 (2023).

63. A. J. Beel, M. Azubel, P.-J. Matteï, R. D. Kornberg, Structure of mitotic chromosomes. Molecular Cell, S1097276521006870 (2021).

64. F. Fatmaoui, P. Carrivain, D. Grewe, B. Jakob, J.-M. Victor, A. Leforestier, M. Eltsov, Cryo-electron tomography and deep learning denoising reveal native chromatin landscapes of interphase nuclei. bioRxiv, 2022.08.16.502515 (2022).

65. Z. Hou, F. Nightingale, Y. Zhu, C. MacGregor-Chatwin, P. Zhang, Structure of native chromatin fibres revealed by Cryo-ET in situ. Nat Commun 14, 6324 (2023).

66. B. Bintu, L. J. Mateo, J.-H. Su, N. A. Sinnott-Armstrong, M. Parker, S. Kinrot, K. Yamaya, A. N. Boettiger, X. Zhuang, Super-resolution chromatin tracing reveals domains and cooperative interactions in single cells. Science 362, eaau1783 (2018).

67. T.-H. S. Hsieh, A. Weiner, B. Lajoie, J. Dekker, N. Friedman, O. J. Rando, Mapping Nucleosome Resolution Chromosome Folding in Yeast by Micro-C. Cell 162, 108–119 (2015).

68. V. I. Risca, S. K. Denny, A. F. Straight, W. J. Greenleaf, Variable chromatin structure revealed by in situ spatially correlated DNA cleavage mapping. Nature 541, 237–241 (2017).

69. S. Cai, D. Böck, M. Pilhofer, L. Gan, The in situ structures of mono-, di-, and trinucleosomes in human heterochromatin. MBoC 29, 2450–2457 (2018).

70. J. K. Chen, T. Liu, S. Cai, W. Ruan, C. T. Ng, J. Shi, U. Surana, L. Gan, Nanoscale analysis of human G1 and metaphase chromatin in situ. bioRxiv, doi: 10.1101/2023.07.31.551204 (2024).

71. P. T. Lowary, J. Widom, New DNA sequence rules for high affinity binding to histone octamer and sequence-directed nucleosome positioning. Journal of Molecular Biology 276, 19–42 (1998).

72. A. Flaus, Principles and practice of nucleosome positioning in vitro. Frontiers in Life Science 5, 5–27 (2011).

73. K. Kelley, A. M. Raczkowski, O. Klykov, P. Jaroenlak, D. Bobe, M. Kopylov, E. T. Eng, G. Bhabha, C. S. Potter, B. Carragher, A. J. Noble, Waffle Method: A general and flexible approach for improving throughput in FIB-milling. Nat Commun 13, 1857 (2022).

74. H. Zhou, J. Hutchings, M. Shiozaki, X. Zhao, L. K. Doolittle, S. Yang, R. Yan, N. Jean, M. Riggi, Z. Yu, E. Villa, M. K. Rosen, Quantitative Spatial Analysis of Chromatin Biomolecular Condensates using Cryo-Electron Tomography. bioRxiv, 2024.12.01.626131 (2024).

75. F. Gordon, K. Luger, J. C. Hansen, The Core Histone N-terminal Tail Domains Function Independently and Additively during Salt-dependent Oligomerization of Nucleosomal Arrays. Journal of Biological Chemistry 280, 33701–33706 (2005).

76. K. Maeshima, R. Rogge, S. Tamura, Y. Joti, T. Hikima, H. Szerlong, C. Krause, J. Herman, E. Seidel, J. DeLuca, T. Ishikawa, J. C. Hansen, Nucleosomal arrays self-assemble into supramolecular globular structures lacking 30-nm fibers. EMBO J 35, 1115–1132 (2016).

77. J. C. Hansen, K. Maeshima, M. J. Hendzel, The solid and liquid states of chromatin. Epigenetics & Chromatin 14, 50 (2021).

78. A. Allahverdi, Q. Chen, N. Korolev, L. Nordenskiöld, Chromatin compaction under mixed salt conditions: Opposite effects of sodium and potassium ions on nucleosome array folding. Sci Rep 5, 8512 (2015).

79. N. Korolev, A. Allahverdi, Y. Yang, Y. Fan, A. P. Lyubartsev, L. Nordenskiöld, Electrostatic Origin of Salt-Induced Nucleosome Array Compaction. Biophysical Journal 99, 1896–1905 (2010).

80. M. Kruithof, F.-T. Chien, A. Routh, C. Logie, D. Rhodes, J. Van Noort, Single-molecule force spectroscopy reveals a highly compliant helical folding for the 30-nm chromatin fiber. Nat Struct Mol Biol 16, 534–540 (2009).

81. J. Widom, Physicochemical studies of the folding of the 100 Å nucleosome filament into the 300 Å filament. Journal of Molecular Biology 190, 411–424 (1986).

82. N. Jentink, C. Purnell, B. Kable, M. T. Swulius, S. A. Grigoryev, Cryo-electron tomography reveals the multiplex anatomy of condensed native chromatin and its unfolding by histone citrullination. Molecular Cell 83, 3236–3252.e7 (2023).

83. M. Zhang, C. Díaz-Celis, J. Liu, J. Tao, P. D. Ashby, C. Bustamante, G. Ren, Angle between DNA linker and nucleosome core particle regulates array compaction revealed by individual-particle cryo-electron tomography. Nat Commun 15, 4395 (2024).

84. L. Chen, M. J. Maristany, S. E. Farr, J. Luo, B. A. Gibson, L. K. Doolittle, J. Rene Espinosa, J. Huertas, S. Redding, R. Collepardo-Guevara, M. K. Rosen, Nucleosome Spacing Can Fine-Tune Higher Order Chromatin Assembly. bioRxiv, 2024.12.23.627571 (2024).

85. A. Golembeski, J. Lequieu, A Molecular View into the Structure and Dynamics of Phase-Separated Chromatin. J. Phys. Chem. B 128, 10593–10603 (2024).

86. T. Schalch, S. Duda, D. F. Sargent, T. J. Richmond, X-ray structure of a tetranucleosome and its implications for the chromatin fibre. Nature 436, 138–141 (2005).

87. F. Song, P. Chen, D. Sun, M. Wang, L. Dong, D. Liang, R.-M. Xu, P. Zhu, G. Li, Cryo-EM Study of the Chromatin Fiber Reveals a Double Helix Twisted by Tetranucleosomal Units. Science 344, 376–380 (2014).

88. S. E. Farr, E. J. Woods, J. A. Joseph, A. Garaizar, R. Collepardo-Guevara, Nucleosome plasticity is a critical element of chromatin liquid– liquid phase separation and multivalent nucleosome interactions. Nat Commun 12, 2883 (2021).

89. G. Arya, T. Schlick, A Tale of Tails: How Histone Tails Mediate Chromatin Compaction in Different Salt and Linker Histone Environments. J. Phys. Chem. A 113, 4045–4059 (2009).

90. Y. Qiu, S. Liu, X. Lin, I. C. Unarta, X. Huang, B. Zhang, Nucleosome condensate and linker DNA alter chromatin folding pathways and rates. bioRxiv, 2024.11.15.623891 (2024).

91. P. J. J. Robinson, W. An, A. Routh, F. Martino, L. Chapman, R. G. Roeder, D. Rhodes, 30 nm Chromatin Fibre Decompaction Requires both H4-K16 Acetylation and Linker Histone Eviction. Journal of Molecular Biology 381, 816–825 (2008).

92. R. Collepardo-Guevara, G. Portella, M. Vendruscolo, D. Frenkel, T. Schlick, M. Orozco, Chromatin Unfolding by Epigenetic Modifications Explained by Dramatic Impairment of Internucleosome Interactions: A Multiscale Computational Study. J. Am. Chem. Soc. 137, 10205–10215 (2015).

93. M. Shogren-Knaak, H. Ishii, J.-M. Sun, M. J. Pazin, J. R. Davie, C. L. Peterson, Histone H4-K16 Acetylation Controls Chromatin Structure and Protein Interactions. Science 311, 844–847 (2006).

94. B. Dorigo, T. Schalch, A. Kulangara, S. Duda, R. R. Schroeder, T. J. Richmond, Nucleosome Arrays Reveal the Two-Start Organization of the Chromatin Fiber. Science 306, 1571–1573 (2004).

95. J. Zhou, J. Y. Fan, D. Rangasamy, D. J. Tremethick, The nucleosome surface regulates chromatin compaction and couples it with transcriptional repression. Nat Struct Mol Biol 14, 1070–1076 (2007).

96. Q. Chen, R. Yang, N. Korolev, C. F. Liu, L. Nordenskiöld, Regulation of Nucleosome Stacking and Chromatin Compaction by the Histone H4 N-Terminal Tail–H2A Acidic Patch Interaction. Journal of Molecular Biology 429, 2075–2092 (2017).

97. T. S. Lewis, V. Sokolova, H. Jung, H. Ng, D. Tan, Structural basis of chromatin regulation by histone variant H2A.Z. Nucleic Acids Research 49, 11379–11391 (2021).

98. W. Alvarado, J. Moller, A. L. Ferguson, J. J. De Pablo, Tetranucleosome Interactions Drive Chromatin Folding. ACS Cent. Sci. 7, 1019–1027 (2021).

99. H. Kenzaki, S. Takada, Linker DNA Length is a Key to Tri-nucleosome Folding. Journal of Molecular Biology 433, 166792 (2021).

100. R. Collepardo-Guevara, T. Schlick, Chromatin fiber polymorphism triggered by variations of DNA linker lengths. Proc. Natl. Acad. Sci. U.S.A. 111, 8061–8066 (2014).

101. O. PerišI\, R. Collepardo-Guevara, T. Schlick, Modeling Studies of Chromatin Fiber Structure as a Function of DNA Linker Length. Journal of Molecular Biology 403, 777–802 (2010).

102. S. Bilokapic, M. Strauss, M. Halic, Histone octamer rearranges to adapt to DNA unwrapping. Nat Struct Mol Biol 25, 101–108 (2018).

103. Y. Takizawa, C.-H. Ho, H. Tachiwana, H. Matsunami, W. Kobayashi, M. Suzuki, Y. Arimura, T. Hori, T. Fukagawa, M. D. Ohi, M. Wolf, H. Kurumizaka, Cryo-EM Structures of Centromeric Tri-nucleosomes Containing a Central CENP-A Nucleosome. Structure 28, 44–53.e4 (2020).

104. M. Dombrowski, M. Engeholm, C. Dienemann, S. Dodonova, P. Cramer, Histone H1 binding to nucleosome arrays depends on linker DNA length and trajectory. Nat Struct Mol Biol 29, 493–501 (2022).

105. S. Kilic, S. Felekyan, O. Doroshenko, I. Boichenko, M. Dimura, H. Vardanyan, L. C. Bryan, G. Arya, C. A. M. Seidel, B. Fierz, Single-molecule FRET reveals multiscale chromatin dynamics modulated by HP1α. Nat Commun 9, 235 (2018).

106. B.-R. Zhou, J. Jiang, R. Ghirlando, D. Norouzi, K. N. Sathish Yadav, H. Feng, R. Wang, P. Zhang, V. Zhurkin, Y. Bai, Revisit of Reconstituted 30-nm Nucleosome Arrays Reveals an Ensemble of Dynamic Structures. Journal of Molecular Biology 430, 3093–3110 (2018).

107. H.-X. Zhou, D. Kota, S. Qin, R. Prasad, Fundamental Aspects of Phase-Separated Biomolecular Condensates. Chem. Rev. 124, 8550–8595 (2024).

108. D. Kota, R. Prasad, H.-X. Zhou, Adenosine Triphosphate Mediates Phase Separation of Disordered Basic Proteins by Bridging Intermolecular Interaction Networks. J. Am. Chem. Soc. 146, 1326–1336 (2024).

109. S. Iida, S. Shinkai, Y. Itoh, S. Tamura, M. T. Kanemaki, S. Onami, K. Maeshima, Single-nucleosome imaging reveals steady-state motion of interphase chromatin in living human cells. Sci. Adv. 8, eabn5626 (2022).

110. I. Y. Wong, M. L. Gardel, D. R. Reichman, E. R. Weeks, M. T. Valentine, A. R. Bausch, D. A. Weitz, Anomalous Diffusion Probes Micro-structure Dynamics of Entangled F-Actin Networks. Phys. Rev. Lett. 92, 178101 (2004).

111. L.-H. Cai, S. Panyukov, M. Rubinstein, Mobility of Nonsticky Nano-particles in Polymer Liquids. Macromolecules 44, 7853–7863 (2011).

112. L.-H. Cai, S. Panyukov, M. Rubinstein, Hopping Diffusion of Nanoparticles in Polymer Matrices. Macromolecules 48, 847–862 (2015).

113. M. Shayegan, R. Tahvildari, K. Metera, L. Kisley, S. W. Michnick, S. R. Leslie, Probing Inhomogeneous Diffusion in the Microenvironments of Phase-Separated Polymers under Confinement. J. Am. Chem. Soc. 141, 7751–7757 (2019).

114. M. L. Gardel, K. E. Kasza, C. P. Brangwynne, J. Liu, D. A. Weitz, “Chapter 19 Mechanical Response of Cytoskeletal Networks” in Methods in Cell Biology (Elsevier, 2008; https://linkinghub.elsevier.com/re-trieve/pii/S0091679X08006195)vol. 89, xpp. 487–519.

115. B. E. Vos, T. M. Muenker, T. Betz, Characterizing intracellular mechanics via optical tweezers-based microrheology. Current Opinion in Cell Biology 88, 102374 (2024).

116. M. Lee, H. C. Moon, H. Jeong, D. W. Kim, H. Y. Park, Y. Shin, Optogenetic control of mRNA condensation reveals an intimate link between condensate material properties and functions. Nat Commun 15, 3216 (2024).

117. Y. Dai, Z. Zhou, W. Yu, Y. Ma, K. Kim, N. Rivera, J. Mohammed, E. Lantelme, H. Hsu-Kim, A. Chilkoti, L. You, Biomolecular condensates regulate cellular electrochemical equilibria. Cell, S0092867424009097 (2024).

118. T. Wu, M. R. King, Y. Qiu, M. Farag, R. V. Pappu, M. D. Lew, Single fluorogen imaging reveals distinct environmental and structural features of biomolecular condensates. Biophysics [Preprint] (2023). 10.1101/2023.01.26.525727.

119. C. Mathieu, R. V. Pappu, J. P. Taylor, Beyond aggregation: Pathological phase transitions in neurodegenerative disease. Science 370, 56–60 (2020).

120. E. M. Hildebrand, J. Dekker, Mechanisms and Functions of Chromosome Compartmentalization. Trends in Biochemical Sciences 45, 385–396 (2020).

121. A. J. Bannister, T. Kouzarides, Regulation of chromatin by histone modifications. Cell Res 21, 381–395 (2011).

122. Q. Szabo, A. Donjon, I. Jerković, G. L. Papadopoulos, T. Cheutin, B. Bonev, E. P. Nora, B. G. Bruneau, F. Bantignies, G. Cavalli, Regulation of single-cell genome organization into TADs and chromatin nanodomains. Nat Genet 52, 1151–1157 (2020).

123. J. Xu, X. Xu, D. Huang, Y. Luo, L. Lin, X. Bai, Y. Zheng, Q. Yang, Y. Cheng, A. Huang, J. Shi, X. Bo, J. Gu, H. Chen, A comprehensive benchmarking with interpretation and operational guidance for the hierarchy of topologically associating domains. Nat Commun 15, 4376 (2024).

124. M. Hu, S. Wang, Chromatin Tracing: Imaging 3D Genome and Nucleome. Trends in Cell Biology 31, 5–8 (2021).

125. S. Wang, J.-H. Su, B. J. Beliveau, B. Bintu, J. R. Moffitt, C. Wu, X. Zhuang, Spatial organization of chromatin domains and compartments in single chromosomes. Science 353, 598–602 (2016).

126. L. Tan, D. Xing, C.-H. Chang, H. Li, X. S. Xie, Three-dimensional genome structures of single diploid human cells. Science 361, 924–928 (2018).

127. T. J. Stevens, D. Lando, S. Basu, L. P. Atkinson, Y. Cao, S. F. Lee, M. Leeb, K. J. Wohlfahrt, W. Boucher, A. O’Shaughnessy-Kirwan, J. Cramard, A. J. Faure, M. Ralser, E. Blanco, L. Morey, M. Sansó, M. G. S. Palayret, B. Lehner, L. Di Croce, A. Wutz, B. Hendrich, D. Klenerman, E. D. Laue, 3D structures of individual mammalian genomes studied by single-cell Hi-C. Nature 544, 59–64 (2017).

128. V. B. Teif, J.-P. Mallm, T. Sharma, D. B. Mark Welch, K. Rippe, R. Eils, J. Langowski, A. L. Olins, D. E. Olins, Nucleosome repositioning during differentiation of a human myeloid leukemia cell line. Nucleus 8, 188–204 (2017).

129. G. Koulouras, A. Panagopoulos, M. A. Rapsomaniki, N. N. Giakoumakis, S. Taraviras, Z. Lygerou, EasyFRAP-web: a web-based tool for the analysis of fluorescence recovery after photobleaching data. Nucleic Acids Research 46, W467–W472 (2018).

130. D. Tegunov, P. Cramer, Real-time cryo-electron microscopy data preprocessing with Warp. Nat Methods 16, 1146–1152 (2019).

131. S. Zheng, G. Wolff, G. Greenan, Z. Chen, F. G. A. Faas, M. Bárcena, A. J. Koster, Y. Cheng, D. A. Agard, AreTomo: An integrated software package for automated marker-free, motion-corrected cryo-electron tomographic alignment and reconstruction. Journal of Structural Biology: X 6, 100068 (2022).

132. Y.-T. Liu, H. Zhang, H. Wang, C.-L. Tao, G.-Q. Bi, Z. H. Zhou, Isotropic reconstruction for electron tomography with deep learning. Nat Commun 13, 6482 (2022).

133. T. Wagner, S. Raunser, The evolution of SPHIRE-crYOLO particle picking and its application in automated cryo-EM processing workflows. Commun Biol 3, 61 (2020).

134. E. Moebel, A. Martinez-Sanchez, L. Lamm, R. D. Righetto, W. Wietrzynski, S. Albert, D. Larivière, E. Fourmentin, S. Pfeffer, J. Ortiz, W. Baumeister, T. Peng, B. D. Engel, C. Kervrann, Deep learning improves macromolecule identification in 3D cellular cryo-electron tomograms. Nat Methods 18, 1386–1394 (2021).

135. J. Zivanov, J. Otón, Z. Ke, A. Von Kügelgen, E. Pyle, K. Qu, D. Morado, D. Castaño-Díez, G. Zanetti, T. A. Bharat, J. A. Briggs, S. H. Scheres, A Bayesian approach to single-particle electron cryo-tomography in RELION-4.0. eLife 11, e83724 (2022).

136. T. A. M. Bharat, C. J. Russo, J. Löwe, L. A. Passmore, S. H. W. Scheres, Advances in Single-Particle Electron Cryomicroscopy Structure Determination applied to Sub-tomogram Averaging. Structure 23, 1743–753 (2015).

137. E. C. Meng, T. D. Goddard, E. F. Pettersen, G. S. Couch, Z. J. Pearson, J. H. Morris, T. E. Ferrin, UCSF ChimeraX: Tools for structure building and analysis. Protein Science 32, e4792 (2023).

138. U. H. Ermel, S. M. Arghittu, A. S. Frangakis, ArtiaX: An electron tomography toolbox for the interactive handling of sub-tomograms in UCSF ChimeraX. Protein Science 31, e4472 (2022).

139. J. Singh, J. M. Thornton, SIRIUS: An automated method for the analysis of the preferred packing arrangements between protein groups. Journal of Molecular Biology 211, 595–615 (1990).

140. D. N. Mastronarde, S. R. Held, Automated tilt series alignment and tomographic reconstruction in IMOD. Journal of Structural Biology 197, 102–113 (2017).

141. D. Farré-Gil, J. P. Arcon, C. A. Laughton, M. Orozco, CGeNArate: a sequence-dependent coarse-grained model of DNA for accurate atomistic MD simulations of kb-long duplexes. Nucleic Acids Research 52, 6791–6801 (2024).

142. A. Jamakovic, P. Van Mieghem, “On the Robustness of Complex Networks by Using the Algebraic Connectivity” in NETWORKING 2008 Ad Hoc and Sensor Networks, Wireless Networks, Next Generation Internet, A. Das, H. K. Pung, F. B. S. Lee, L. W. C. Wong, Eds. (Springer Berlin Heidelberg, Berlin, Heidelberg, 2008; http://link.springer.com/10.1007/978-3-540-79549-0_16)vol. 4982 of Lecture Notes in Computer Science, pp. 183–194.

143. J. Schindelin, I. Arganda-Carreras, E. Frise, V. Kaynig, M. Longair, T. Pietzsch, S. Preibisch, C. Rueden, S. Saalfeld, B. Schmid, J.-Y. Tinevez, D. J. White, V. Hartenstein, K. Eliceiri, P. Tomancak, A. Cardona, Fiji: an open-source platform for biological-image analysis. Nat Methods 9, 676–682 (2012).

144. T. Kuhn, J. Hettich, R. Davtyan, J. C. M. Gebhardt, Single molecule tracking and analysis framework including theory-predicted parameter settings. Sci Rep 11, 9465 (2021).

145. J. C. M. Gebhardt, D. M. Suter, R. Roy, Z. W. Zhao, A. R. Chapman, S. Basu, T. Maniatis, X. S. Xie, Single-molecule imaging of transcription factor binding to DNA in live mammalian cells. Nat Methods 10, 421–426 (2013).

146. S. Ide, S. Tamura, K. Maeshima, Chromatin behavior in living cells: Lessons from single-nucleosome imaging and tracking. BioEssays 44, 2200043 (2022).

147. T. G. Mason, Estimating the viscoelastic moduli of complex fluids using the generalized Stokes–Einstein equation. Rheologica Acta 39, 371–378 (2000).

148. A. Routh, S. Sandin, D. Rhodes, Nucleosome repeat length and linker histone stoichiometry determine chromatin fiber structure. Proc. Natl. Acad. Sci. U.S.A. 105, 8872–8877 (2008).

149. T. Brouwer, C. Pham, A. Kaczmarczyk, W.-J. de Voogd, M. Botto, P. Vizjak, F. Mueller-Planitz, J. van Noort, A critical role for linker DNA in higher-order folding of chromatin fibers. Nucleic Acids Research 49, 2537–2551 (2021).

150. C. Tse, J. C. Hansen, Hybrid Trypsinized Nucleosomal Arrays: Identification of Multiple Functional Roles of the H2A/H2B and H3/H4 N-Termini in Chromatin Fiber Compaction. Biochemistry 36, 11381–11388 (1997).

151. S. C. Moore, J. Ausió, Major Role of the Histones H3-H4 in the Folding of the Chromatin Fiber. Biochemical and Biophysical Research Communications 230, 136–139 (1997).

152. C. Zheng, X. Lu, J. C. Hansen, J. J. Hayes, Salt-dependent Intra- and Internucleosomal Interactions of the H3 Tail Domain in a Model Oligonucleosomal Array. Journal of Biological Chemistry 280, 33552–33557 (2005).

153. P.-Y. Kan, X. Lu, J. C. Hansen, J. J. Hayes, The H3 Tail Domain Participates in Multiple Interactions during Folding and Self-Association of Nucleosome Arrays. Molecular and Cellular Biology 27, 2084–2091 (2007).

154. P.-Y. Kan, T. L. Caterino, J. J. Hayes, The H4 Tail Domain Participates in Intra- and Internucleosome Interactions with Protein and DNA during Folding and Oligomerization of Nucleosome Arrays. Molecular and Cellular Biology 29, 538–546 (2009).

155. S. Pepenella, K. J. Murphy, J. J. Hayes, A Distinct Switch in Interactions of the Histone H4 Tail Domain upon Salt-dependent Folding of Nucleosome Arrays. Journal of Biological Chemistry 289, 27342–27351 (2014).

156. B. Dorigo, T. Schalch, K. Bystricky, T. J. Richmond, Chromatin Fiber Folding: Requirement for the Histone H4 N-terminal Tail. Journal of Molecular Biology 327, 85–96 (2003).

157. B. R. Macadangdang, A. Oberai, T. Spektor, O. A. Campos, F. Sheng, M. F. Carey, M. Vogelauer, S. K. Kurdistani, Evolution of histone 2A for chromatin compaction in eukaryotes. eLife 3, e02792 (2014).

158. A. Allahverdi, R. Yang, N. Korolev, Y. Fan, C. A. Davey, C.-F. Liu, L. Nordenskiöld, The effects of histone H4 tail acetylations on cation-induced chromatin folding and self-association. Nucleic Acids Research 39, 1680–1691 (2011).

159. X. Wang, J. J. Hayes, Acetylation Mimics within Individual Core Histone Tail Domains Indicate Distinct Roles in Regulating the Stability of Higher-Order Chromatin Structure. Molecular and Cellular Biology 28, 227–236 (2008).

160. D. Sinha, M. A. Shogren-Knaak, Role of Direct Interactions between the Histone H4 Tail and the H2A Core in Long Range Nucleosome Contacts. Journal of Biological Chemistry 285, 16572–16581 (2010).

