## Supporting Information for "Multi-scale structure of chromatin condensates rationalizes phase separation and material properties"

bioRχiv

**This PDF file includes:**

Materials and Methods
Supplementary Text
Figs. S1 to S5
Tables S1 to S4
References

### Materials and Methods

#### Protein and Nucleosome Arrays, Native Chromatin

Nucleosome arrays (*33*), HeLa nuclei, and NIH3T3 cells (*74*) were prepared as described previously.

#### Chromatin Phase Separation

Chromatin condensates were induced by combining 6 μM of either 25 bp or 30 bp nucleosome arrays with a 2X phase separation buffer (40 mM Tris-OAc pH 7.5, 300 mM KOAc, 5% glycerol, 0.2 mM EGTA). These condensates were subsequently used in a series of experiments, including FRAP, droplet fusion, cryo-ET, trypsin digestion, single-molecule tracking, and microrheology.

#### Phase Diagram with Turbidity Measurement

The extent of phase separation was assessed by measuring absorbance at 345 nm using a NanoDrop One UV-Vis spectrophotometer (Thermo Scientific). Nucleosome arrays were diluted to the desired concentration using a buffer containing 20 mM Tris-OAc pH 7.5, 0.1 mM EGTA, and mixed with 2X phase separation buffer at varying KOAc concentrations. Each sample was measured three times, and the results were averaged.

#### FRAP of Chromatin Condensate

FRAP experiments were performed using a Leica SP8 confocal fluorescence microscope equipped with a 561 nm laser for photobleaching. Prior to photobleaching, three frames were captured to establish a baseline. A region of interest (ROI) within the chromatin condensate was selectively bleached with a high-intensity laser for 5 seconds. Fluorescence recovery within the bleached ROI was monitored at 5-second intervals over a total period of 300 seconds.

Images were captured using PMT detectors paired with a 60x oil immersion objective. Fluorescence intensity within the ROI was quantified before and after bleaching using Leica LAS-X software. Recovery curves were plotted to determine the mobile fraction and the half-time of fluorescence recovery. Data analysis was conducted using easyFRAP (129), and the results were used to calculate the kinetics of chromatin mobility. The FRAP curves were fitted using double-scale normalization.

#### Imaging and Analyzing Droplet Fusion

Real-time imaging of droplet fusion was conducted using a Leica DMI6000 microscope, equipped with a Yokogawa CSU-X1 spinning disk confocal scanner unit and a Hamamatsu ImagEMX2 EM-CCD camera. A Leica 40x oil immersion objective was employed for imaging. Time-lapse sequences were recorded at 100 ms per frame.
Images were acquired using Metamorph (Biovision) software. The time-lapse images were analyzed with FIJI/ImageJ software to track droplet fusion dynamics. Fusion events were identified manually. To determine the time constant ($\tau$) for each condensate fusion event, the change in aspect ratio (AR) over time was measured, fitting the values to an exponential decay model: $AR = 1 + (ARinit - 1) * \exp(-t/\tau)$, where ARinit represents the initial aspect ratio at the onset of fusion.

#### Trypsin Digestion and Turbidity Measurement

Chromatin condensates (22 μL), prepared as described above, were mixed with 3 μL of trypsin (4 ng/μL in 25 mM Tris-HCl, pH 7.5) in an mPEG-Silane pretreated microscopic plate (35). The plate was immediately placed in a plate reader (Tecan Spark) to monitor turbidity changes by measuring absorption at a wavelength of 345 nm until the signal reached a plateau. Turbidity values were normalized within each well by subtracting the plateau value from the readings throughout the digestion process. The measurements were performed in three independent wells, and the results were averaged. Standard deviations are shown to indicate variability.

#### Cryo-ET Sample Preparation and Data Collection

##### Chromatin samples in low salt buffer

To prepare chromatin samples under low salt conditions, Lacey carbon grids (300 mesh, Ted Pella Inc) were glow discharged using an easiGlow (Pelco) at 30 mA for 30 seconds. A Vitrobot Mark IV (Thermo Fisher) was pre-cooled to 4°C and set to 100% humidity. A 3 μL aliquot of nucleosome array solution (6 μM in 20 mM Tris-Cl, 0.1 mM EGTA) was applied to the grid, blotted with a force setting of 0 for 4 seconds, and plunge frozen in liquid ethane.

##### Chromatin sample in high salt buffer

A 4 μL aliquot of phase separated nucleosome array solution (6 μM in 20 mM Tris-Cl, 150 mM KOAc, 0.1 mM EGTA) was vitrified using the waffle method (73) on a Wohlwend Compact 03 high-pressure freezer as previously described (74). After freezing, grids were analyzed by cryo-confocal imaging (Leica EM Cryo-CLEM system equipped with a 50x/dry objective (NA=0.9)) to identify regions containing condensates, then transferred to an Aquilos II cryo-FIB mill (Thermo Fisher). Lamellae (60-200 nm thick) were randomly generated in these regions using notch-milling procedures, and condensates were identified using the SEM mode of the instrument, as previously described (73, 74).

#### Cryo-ET Data Collection

For chromatin samples with a 25 bp linker length in low salt conditions, tilt-series were collected using a 300 kV Titan Krios G1 (Thermo Fisher Scientific) equipped with a high-brightness field emission gun (xFEG), a spherical aberration corrector, a BioQuantum energy filter, and a post-GIF K3 camera (Gatan Inc.). Tilt series were acquired using a dose symmetric scheme over a tilt range from -60° to +60° with a 3-degree increment. For each tilt angle, a movie was recorded at a nominal magnification of 33,000x, with a calibrated physical pixel size of 0.206 nm in CDS (correlated double sampling) counted mode. A 20 eV energy slit was used and the total electron dose was limited to 162 electrons/Å$^2$. A Volta phase plate was employed, with a targeted defocus set to -0.5 μm (table S5).

For chromatin samples with a 30 bp linker length in low salt conditions, tilt-series were collected using a Titan Krios G3i (Thermo Fisher) equipped with a cold field emission gun, a Selectris X energy filter, and a Falcon 4i camera. Images in each tilt series were acquired using a dose symmetric scheme over a tilt range from -45° to +45° with a 3-degree increment. For each tilt angle, a movie was recorded at a nominal magnification of 53,000x with a calibrated pixel size of 0.23 nm.. A 5eV energy slit was used and the total electron dose was limited to 120 electrons/Å$^2$ (table S5).

For chromatin condensates in high salt conditions, data collection was performed using the Krios G3i as described above. The tilt series spanned angles from -48° to +60° with 2° increments per tilt. Images were recorded at a physical pixel size of 0.1516 nm at the specimen level. The defocus values ranged from -3 μm to -4.5 μm, and the total electron dose was limited to 178 electrons/Å$^2$ (table S5, (74)).

### Cryo-ET Data Analysis

The tilt-series data were pre-processed using Warp (130) for motion correction and CTF estimation, then subsequently aligned with AreTomo (131). The alignments were then used in Warp to reconstruct two half-sets tomogram with a pixel size of 8 Å and denoised sequentially (74) using Warp (130) and IsoNet (132).

For tomograms of chromatin under low salt conditions, nucleosome particles were picked using crYOLO (133). A subset of tomogram slices was manually annotated for training purposes. After particle picking, the tomograms were manually inspected to remove junk particles and particles closer than ~30 nm to the air-water interface.

Tomograms of chromatin condensates were initially segmented using DeepFinder (134), and the centers of particles were localized with the MeanShift algorithm. The particles with estimated positions were then processed through our custom algorithm, Context-Aware Template Matching (CATM), to determine nucleosome orientation and refine particle centers(74). Briefly, the particles were extracted from the tomogram and subjected to local template matching using a nucleosome structure low-pass filtered at 25 Å. For each particle, multiple positions and orientations, along with their corresponding cross-correlation coefficients, were recorded. This information was then used to optimize particle assignments and eliminate clashes. Once nucleosome orientations and positions were accurately assigned, sub-tomograms were generated in Warp and further refined in Relion (135, 136) to yield the final positions of all nucleosomes.

### Tri-Nucleosome Model Building

To build tri-nucleosome models (fig. S1C), individual nucleosomes were first aligned to reconstruct an average mono-nucleosome. The refinement box size was then expanded to align adjacent nucleosomes, which were identified at either DNA end of the central nucleosome in separate classes. The di-nucleosome structures, with adjacent nucleosomes at either end, were then aligned based on the central nucleosome to create a tri-nucleosome structure for visualization (note that all geometric analyses were performed using individual nucleosomes, not these averages, which are purely for visualization). Due to the limited number of particles available, we were unable to achieve high-quality sub-tomogram averaging for the 30 bp structure in high-salt dilute phase conditions. Therefore, we used the tri-nucleosome structure of the 30 bp chromatin in the high-salt condensate phase as a representative in Fig. 2F, as the conformations under these two conditions are very similar (fig. S3C-D, G-I).

### Tracing of Individual Nucleosome Arrays

The tracing process begins by positioning the assigned nucleosomes into the tomograms using ChimeraX (137) with the Artix plugin (138) for visual inspection. Erroneous assignments were manually removed, and the in-plane rotation of nucleosomes was adjusted based on their shape and linker DNA density. Connectivity between nucleosomes within the array was manually assigned. The traced nucleosome positions and orientations were further refined in CATM (74) to locally optimize their orientations and resolve particle clashes.

The traced nucleosome arrays were then used to quantify nucleosome conformations. The distance between nucleosome N and N+2 was calculated using Euclidean distance (D) between their centers of mass, and the dihedral angles between adjacent (a) or alternating (para) nucleosomes were determined by calculating the angles between their planes. Dihedral angles were calculated using vectors perpendicular to the nucleosome planes. To calculate a, for the 25 bp nucleosome array, adjacent nucleosome vectors were oriented oppositely, reflecting the 2.5 turns of DNA between them. The angle between these vectors was measured and scaled to a range of 180°–270°, where 180° corresponds to parallel nucleosome planes and 270° to perpendicular planes. For the 30 bp array, the vectors were aligned due to the 3 turns of DNA, and the angle between them was calculated and scaled to 0°–90°, with 0° indicating parallel planes and 90° indicating perpendicular planes. To calculate para, the vectors were aligned, and the resulting angle was similarly scaled to 0°–90°.
Classification of Mono, Di, and Tri-Nucleosome Conformations in Chromatin Condensates
To assess mono-nucleosome structural variability, 3D classification was performed using Relion. Subtomograms were sampled at a pixel size of 8 Å with 64x64x64 voxels. Nucleosomes were classified into six classes over ten iterations in Relion using a 200 Å diameter spherical mask. The resulting nucleosome structures were aligned based on one linker DNA to observe structural variability in the other linker DNA.

To analyze 25 bp chromatin structural variability at the di-nucleosome level, nucleosomes were classified into eight classes. This analysis produced 35.8% of nucleosomes with both one adjacent nucleosome and the linker DNA well resolved; these nucleosomes were categorized into three different classes. To analogously explore the conformational diversity of multi-nucleosome groups in 30 bp chromatin, nucleosomes were classified, producing 56.5% with well-resolved linker DNA and two adjacent nucleosomes. These well-resolved particles were further organized into two different nucleosome stacking patterns: 20.6% of particles exhibited an offset between stacked nucleosomes, while 79.4% of particles displayed perfect face-to-face stacking (Fig. 4H).

### Nucleosome Pair Orientations

In our coordinate system, the origin is set at the centroid of the reference nucleosome, which is approximated as having 2-fold symmetry about the dyad axis (i.e., composed of palindromic DNA). The Z-axis is defined as the direction along which the nucleosome projection is maximum (i.e. perpendicular to the nucleosome plane). The X-axis extends from the centroid to the nucleosome dyad, and the Y-axis is orthogonal to the X-Z plane. For each nucleosome identified through template matching, we calculated the angle between its Z-axis and either the beam direction or the two directions perpendicular to the beam. A random arrangement of nucleosomes will exhibit a sinusoidal distribution of each angle(139).

The relative orientation of two nucleosomes was classified into three categories: face-to-face, side-to-side, and face-to-side (Fig. 5F and 7I), similar to definitions described by Farr and colleagues (88). Classification was based on vectors perpendicular to the nucleosome planes (i.e. along the Z-axes) through their centroids according to the following algorithm:
For nucleosomes $i$ and $j$, the angles $\theta$ and $\psi$ are defined according to:
$\hat{z}i \cdot \hat{z}j = \cos\theta$
$\hat{z}i \cdot \hat{r} = \cos\psi i$
$\hat{z}j \cdot \hat{r} = \cos\psi j$

where $\hat{z}_i$ and $\hat{z}_j$ are the unit vectors of nucleosomes i and j, and $\hat{r}$ is the unit vector pointing from the centroid of nucleosome i to the centroid of nucleosome j. Thus, $\theta$ denotes the angle between the two unit nucleosome vectors (used in Figures 5B and 7H), and $\psi$ denotes the angle between a nucleosome unit vector and the line between the nucleosome centroids.

```
if θ < 45° or θ > 135°:                                  {nucleosomes ~parallel)
if ψi < 60° or ψi < 120° or ψj < 60° or ψj > 120°:       {one nucleosome above the other}
  face-to-face
else:                                                    {nucleosomes beside each other}
    side-to-side
else:                                                    {nucleosomes ~perpendicular)
  if ψi > 45 and ψi < 135 and ψj > 45 and ψj < 135:      {nucleosome edges close to each other}
    side-to-side
  else:                                                  {perpendicular packing}
     face-to-side
```

Only nucleosome pairs whose centroids were closer than 12 nm were considered in the analysis of relative orientation.

### Radial Distribution Analysis

To analyze the radial distribution of nucleosomes within the condensates, a spherical region with 60 nm diameter centered on each nucleosome was used to estimate the local nucleosome concentration. To avoid edge artifacts, the 500 nucleosomes with the highest local concentrations were used in the analysis to ensure they were fully within the condensates. For each nucleosome, a series of concentric shells of 1 nm thickness with outer diameters incrementing by 1 nm up to 60 nm was generated. The number of particles within each spherical shell was calculated and normalized against the average nucleosome concentration and the volume of the shell. The distribution was then averaged across all nucleosomes.

### Radius of Gyration (Rg) Calculation

To calculate the radius of gyration for nucleosome arrays, we first identified the center of mass by averaging the coordinates of each nucleosome position. Next, we computed the squared distances of each nucleosome from this center of mass. The radius of gyration was obtained as the square root of the mean of these squared distances. Only arrays with more than 4 nucleosomes were considered, and Rg was normalized by scaling it with the square root of the ratio between the reference count (12) and the actual number of nucleosomes.

### Segmentation of Chromatin Domains in Native Chromatin Tomograms

To examine chromatin segregation within tomograms of Hela nuclei and NIH3T3 cell nuclei, we used IMOD (140) to manually delineate the segregated domains. The assigned nucleosomes were then mapped back to their respective domains and visualized in ChimeraX (137) using the ArtiaX plugin (138).
Multiscale Modelling of Chromatin Fibers in the Dilute and Condensed phases
To represent chromatin fibers, we use our multiscale chromatin model (88), a bottom-up approach that combines atomistic information of nucleosomes, DNA, and proteins, with two levels of coarse-graining: chemical-specific and minimal. This model enables us to explore how the chemical composition of chromatin influences its structure and propensity to phase separate. We adopted a multiscale strategy to take advantage of both (1) the precision of atomistic models, which can reveal how chemical modifications alter the local behavior of proteins and DNA, describe protein-chromatin binding, and explain how DNA sequence affects mechanical properties, and (2) the efficiency of coarse-grained models, which significantly reduce system dimensionality—for example, representing a 100 kb chromatin region (approximately 10 million atoms plus solvent) with just ~15K beads. These two coarse-grained models are described below.

### Chemical-Specific Coarse-grained Model

The chemical specific chromatin coarse-grained model explicitly represents each amino acid in the histone proteins as a bead centered on its alpha-carbon. Beads corresponding to lysine, arginine, aspartic acid, glutamic acid, and histidine carry the full charge of their atomistic counterparts at pH ~7. Each amino acid bead has a relative hydrophobicity and diameter values derived from atomistic simulations and experimental data (88). The histone protein core is modeled using an elastic network that maintains the secondary structure of histones in the 1KX5 crystal structure. Histone tails are treated as flexible polymers, with bonds between consecutive residues maintained by a stiff harmonic potential, without penalizing for bending or torsion.

Screened electrostatic interactions between all non-bonded beads are approximated using the Debye-Hückel model. Non-ionic associations are modeled with a Lennard–Jones potential. All model parameters and energy functions are detailed in the previous studies(88, 141).

### Minimal Coarse-Grained Chromatin Model.

Our minimal chromatin model (88) represents the histone core as a single bead, modeled as an ellipsoid with radii of $28 \times 28 \times 20$ Å. Linker and nucleosomal DNA are represented by finite-size orientable spheres (an ellipsoid of $12 \times 12 \times 12$ Å), with one sphere per 5 base-pair segment. To capture the mechanical properties of DNA, we developed a minimal "Rigid-Base-Pair-like" model with a resolution of 5 base pairs per bead. The minimal helical parameters were optimized based on chemically specific coarse-grained simulations of 200 bp DNA strands (88). Nucleosome–nucleosome and nucleosome–DNA interactions are modeled using a set of orientation-dependent potentials, fitted to reproduce the internucleosome potentials of mean force obtained from our chemically specific coarse-grained chromatin model. All model parameters and energy functions are detailed in our pervious study(88).

### Reconstruction of Chromatin Fibers in the Dilute Phase

Flat-bottomed harmonic restraints were used to impose the inter-nucleosomal distances obtained from cryo-EM tracing onto simulated nucleosome arrays using our GPU-accelerated chemical-specific coarse-grained chromatin model. The restraints were implemented in OpenMM using a CustomCentroidBondForce with the following expression:
with r being the centroid distance between each 2 nucleosomes. A value of 5 kJ/mol/nm2 was used for k. The value of tolerance used for a given bond incorporated the cross-correlation coefficient (CCC) obtained from the cryo-EM maps, with the formula 1 – CCCaverage being used to calculate the final tolerance, where CCCaverage is the average CCC of the two nucleosomes the constraint is applied to.

$$V(r) = \begin{cases} 0.5\,k \cdot (r - (r_{\text{eq}} + \text{tolerance}))^2 & \text{if } r > r_{\text{eq}} + \text{tolerance}, \\ 0.5\,k \cdot (r - (r_{\text{eq}} - \text{tolerance}))^2 & \text{if } r < r_{\text{eq}} - \text{tolerance}, \\ 0 & \text{otherwise.} \end{cases}$$

The tolerance was gradually decreased from 2(1 – CCCaverage) to the final value in five sets of 50 ns simulations. All initial steps were performed at a salt concentration of 60 mM, before the simulation was ran for 500 ns at its target salt concentration (100 mM for high salt and 25 mM for low salt). The last 400 ns were retained for analysis.

### Reconstruction of Chromatin Fibers inside Condensates

In principle we could have used the approach described above for dilute phase fibers (Fig. 3) to analyze condensate fibers surrounded by their neighboring nucleosomes (Fig. 5c,d). However, we were concerned that because the surrounding nucleosomes could not, in most cases, be connected to one another, tail behaviors of the fibers might be in error. To address this concern we developed an alternative, multiscale approach that leverages coarse-grained models at two resolutions (88) (fig. S4a). As described in Figure S4, coarse-grained simulations using the low resolution, minimal model of 25 bp and 30 bp chromatin condensates produced collections of molecular structures whose distributions of Rg values contained those of the molecules that we could trace in the experimental tomograms. Moreover, we recently showed that these simulations quantitatively recapitulate the experimentally determined phase separation thresholds of a series of chromatin molecules (84). Thus, here we first used direct coexistence simulations using the minimal model, to generate approximately 10,000 uncorrelated equilibrium configurations of chromatin condensates, each containing 125 interacting 12-nucleosome chromatin fibers at high salt, with either 25 or 30 bp linkers (fig. S4a, iv; note that these same coarse grained simulation trajectories were also reported in (84)). Then for each of the ten fibers that could be traced in the cryo-ET tomograms for each chromatin type, we searched the full simulation ensemble for the best match (among ~1,250,000 individual fibers). Scoring was based on the root mean square deviation (RMSD) between the geometric centers of the nucleosomes in the cryo-ET and simulated fibers (fig. S4a i-ii, and iii, respectively). The top-scoring (lowest RMSD) fiber was tracked back to its simulated condensate (fig. S4a iv) and extracted along with its full interaction cluster—i.e., all neighboring fibers that contacted it (fig. S4a v).

Subsequently, this cluster was backmapped from the minimal coarse-grained model to our higher resolution chemically specific model, enabling its reconstruction at the amino acid and nucleotide level (fig. S4a vi). To perform this backmapping, we used a shorter, 15 ns version of the protocol for dilute reconstructions with tolerance set to 0, as the coordinates obtained from the minimal model have no associated uncertainty. The reconstructed models were then added one by one to assemble the full interaction cluster, with harmonic restraints applied to constrain the inter-nucleosome distances, maintaining the array conformation. A second CustomCentroidBondForce, with expression:

$$V(r) = k'((x - x_0)^2 + (y - y_0)^2 + (z - z_0)^2)$$

where x.y and z are the coordinates of a nucleosome centroid, and x0, y0 and z0 are the target coordinates of that nucleosome centroid, was then used to pull each newly added array to its correct position, with k' gradually ramping from 0 to 2.5 kJ/mol/nm$^2$ over a period of 2.5 ns each time a new array is added. As before, the entire reconstruction procedure was carried out at a salt concentration of 60 mM, before switching to the final salt concentration of 100 mM once all arrays were in place. The final configuration was then simulated for 500 ns with all restraints in place, with the final 400 ns retained for analysis.

### Histone-tail Contact Analysis

For the analysis of tail-mediated interactions of chromatin arrays in both the dilute phase (low and high salt) and inside the condensates, we built chemically specific coarse-grained models of the chromatin arrays based on the cryo-ET data and performed molecular dynamics simulations as detailed above. For each experimental condition (25 bp low salt dilute phase, 25 bp high salt dilute phase, 30 bp low salt dilute phase, 30 bp high salt dilute phase, 25 bp high salt condensate, 30 bp high salt condensate), we ran independent simulations for each of the available experimentally resolved chromatin arrays (Supplementary Table S1). In each case, we analyzed one simulation frame (i.e., the coordinates of all beads at a fix time-point in the simulation) for every ns within the last 400 ns of the simulation trajectories (post-equilibration), for a total of 400 frames.

For each simulation frame, we computed the following interaction contact matrices, I, for all possible pairs of beads (regardless of whether they are histone or DNA beads), $i$ and $j$, in any nucleosome in the array:

$$I_{ij} = \begin{cases} 1, & \text{if beads } i \text{ and } j \text{ are "in contact"} \\ 0, & \text{otherwise} \end{cases}$$

Here, “in contact” is true whenever the distance between beads i and j is less than a set cutoff distance. The cut-off distance was set to the average van der Waals radius of the two beads plus 0.1 nm.

To plot the histone-tail mediated contacts, we focus on the rows of the matrix corresponding to histone beads. For each histone bead, i, the total number of contacts is computed by adding up its contacts with all other beads (DNA and histone beads) that meet different criteria (labelled “x” below), depending on the type of contact of interest:

$$M^{x}(i) = \sum_{j \text{ in x}}^{L} I_{ij}$$

Here, $M$ is the total number of beads in the array per frame. For the “inter-nucleosome contacts” shown in Figures 3 and 5, the condition $j$ in x is set to count contacts only if they occur among beads of different nucleosomes. For the “inter-array contacts” in Figure 5, $j$ in x is set to count contacts only if they occur among beads of different nucleosomes and different fibers. For the “inter-nucleosome contacts” in Figure 5, $j$ in x is set to count contacts only if they occur among beads of different nucleosomes and the same fibers. For the intra-nucleosome contacts shown in Figure S2, $j$ in x is set to count contacts only if they occur beads of the same nucleosome.

The value of $M^x(i)$ is then added over for all beads that correspond to the same histone bead residue within all the different nucleosomes (e.g. for all beads corresponding to the H4K16 residue across nucleosomes 1, 2, ... N)

$$C^x(\text{histone residue}) = \frac{1}{a} \sum_{i\,=\text{histone residue}}^{L} M^x(i)$$

Here $a$ is a normalization constant. $C^x$ is recalculated for each analysis frame, and all its values area summed. Finally, the accumulated values of $C^x$ are normalized by $2Nt$, where N is the number of nucleosomes in the array, 2 considers the two copies of each tail, and $t$ is the total number of frames used for analysis. The procedure is repeated for all the different simulations corresponding to the same experimental condition. The plots of contacts per residue display the values of $C^x(i)$ averaged across these different simulations.

For the plots where contacts are accumulated per tail (Fig. 3E and Fig. 5I), an additional sum is performed to add up the contacts of all the beads $S$ of each tail:
$T^{\{H4,H3,H2A(N),H2B\}} = \sum_{i \text{ in } \{\text{H4,H3,H2A(N),H2B}\}}^{S} C^x(\text{histone residue})$.

### Chromatin Network Analysis

We used minimal model coarse-grained simulations to analyze the chromatin interaction network within the condensates. For each chromatin type, we performed a 5-10 microseconds simulation and selected 12 uncorrelated configurations for analysis. For each configuration, nucleosomes from every array were identified along with their neighboring nucleosomes. The interactions were classified according to face-to-face, face-to-side, and side-to-side geometries as described above. Based on the potential of mean force (PMF) for di-nucleosome interactions (84), nucleosomes within 80 Å, 90 Å, or 120 Å were considered to be interacting through the three geometries, respectively. For each interaction, an energy of 8 kT, 7.5 kT, or 1.5 kT, respectively, was assigned to the corresponding nucleosome pair. The interaction energy for each pair of arrays was then assigned to the sum of their nucleosome-nucleosome contact energies.

To construct a geometric graph network, each array was represented as a node. Array-array interactions with energies greater than or equal to 7.5 kT then established an edge between the corresponding nodes. The interaction energy was used as the weight for the edge. We evaluated connectivity of the network through the normalized algebraic connectivity. Algebraic connectivity, also known as the Fiedler value, refers to the second smallest eigenvalue of the Laplacian matrix of a graph, which represents the overall connectivity and robustness of the network (142). The algebraic connectivity of the chromatin condensate graphs was normalized using a 3D lattice graph, where the degree of connections matched the maximum degree observed in the chromatin condensate. Normalized algebraic connectivity values for the 12 independent condensate configurations were averaged to describe each system.

To assess network stability, 25% of the interactions were turned off at equilibrium, and the simulation was continued. The chromatin network gradually disassembled, resembling the behavior observed in trypsin digestion experiments. Graph networks were constructed throughout the dissolution process, and the size of the largest giant component was measured.
Single-Molecule Tracking and Analysis
Single-molecule imaging was performed using a DeltaVision OMX SR system, equipped with a 60 × 1.49 NA TIRF oil immersion objective and ring-TIRF system. Imaging was conducted in epi/TIRF mode using CMOS cameras set to 2 × 2 binning and a beam condenser to enhance contrast. For single-molecule tracking of nucleosomal arrays, burst mode was used at 50 Hz imaging rate.

Movies were initially processed using Fiji/ImageJ (143) for background subtraction. The first 500 frames of each movie were discarded to ensure detection of only single nucleosome arrays. Movies were analyzed using TrackIt software (144) . Spots were detected using a threshold factor of 2 and tracked with a tracking radius of 0.8 pixel, a minimal track length of 10, gap frames of 3 and a minimal track length before gap frame of 5.

The tracked data were analyzed via jump distance analysis to determine an effective diffusion coefficient, Deff. In this analysis cumulative distributions of squared jump distances were fit with two exponential components corresponding to two effective diffusion coefficients D1, 2 with amplitudes A1, 2, respectively (145):

$$f_2(X) = A_1\left(1 - e^{\frac{-X}{D1}}\right) + (1 - A_1)\left(e^{\frac{-C1}{D2}} - e^{\frac{-X}{D2}}\right) \Big/ \left(e^{\frac{-C1}{D2}} - e^{\frac{-C2}{D2}}\right)$$

Where X=(x2+y2)/(4τ) with the camera frame rate τ. Functions are normalized to account for the lower and upper limit of jump distances, C1=0 and C2=rmax, where rmax is the tracking radius. Deff was defined as:

$$D_{eff} = \sum_i D_i \times A_i$$

Deff is used as a measure of the overall mobility of the tracked nucleosome arrays. To assess the robustness of the data and for further error estimation, resampling was performed. Fifty percent of jumps from the pooled distribution were randomly drawn 20 times, and each distribution was fitted individually. The average value over all resampling was shown, with the error bar indicating the standard deviation of the resampling values.

To assess the asymmetry of array movement within the condensates, we initially transformed the sequential points {(x0, y0), (x1, y1), ⋯, (xn, yn), ⋯} representing a trajectory in the xy plane into a series of displacement vectors Δrn = (xn+1 − xn, yn+1 − yn)t. We then computed the angle between each pair of vectors Δrn and Δrn+1. The angles were normalized to $2\pi$ for visualization, with color representing the angles and values corresponding to the probability density. The Asymmetry Coefficient (AC) was determined as the log2 of the ratio between the frequencies of forward (FWD) angles (−30° to +30°) and backward (BWD) angles (150° to 210°) (146). A negative AC value indicates a deviation from a uniform distribution, with backward angles being predominant.

$$AC = \text{Log2}\left(\frac{FWD}{BWD}\right)$$

### Video Particle Tracking

Samples were prepared in 384-well glass bottom microwell plates (Brooks Life Science Systems: MGB101-1-2-LK-L). Prior to use, the plates were cleaned with 5% Hellmanex III (Hëlma Analytics), etched with 1 M KOH, and siliconized with Sigmacote (Sigma-Aldrich). On the day of the experiment, individual wells were blocked with 1% bovine serum albumin, then rinsed thoroughly with MilliQ-water. Chromatin samples were added (final concentration of 1 uM) in 25 mM Tris-Acetate pH 7.5, 100 mM KOAc, 5% glycerol, along with 175 nm-diameter carboxylate-modified fluorescent beads (<1500 beads/uL, Invitrogen: P7220). Samples were incubated inside the temperature-controlled microscope chamber at 30 °C for at least 1 hour before imaging.

Images were captured on a Lecia DMI6000 B microscope base with a Yokogawa CSU-X1 spinning disk confocal scanner unit and a 405/488/561/647 nm Laser Quad Band Set filter cube (Chroma) with a plan apo 63 or 100 × 1.40 NA oil immersion objective. Images were acquired using a Hamamatsu ImagEMX2 EM-CCD camera at 15 ms/frame using the stream acquisition function in Metamorph (Biovision) software. A total of 10,000-100,000 frames (2.5-25 min) were acquired for each acquisition, up to 6 acquisitions were made per sample per session, and at least 5 independent sessions were carried out per sample for reproducibility.

Particle tracking and calculation of mean squared displacement (MSD) was performed using MATLAB codes by Daniel Blair and Eric Dufresne (https://site.physics.georgetown.edu/matlab/code.html). Average MSD was calculated from ~10,000 individual trajectories, and smoothed using a moving average with span < 10 % of total number of frames. Elastic (G') and viscous (G") moduli as a function of frequency (ω) were calculated using the generalized Stokes-Einstein relation (GSER) as described by Mason TG (147) using MATLAB codes by Andrew Sun (https://github.com/andrewx101/track_analysis/releases/tag/v2.05). Viscosity ($\eta$) was calculated ($\eta$ = G"/ω) and plotted against frequency. From the viscosity plot, the mean value of the plateau at low frequency was used to estimate the zero-shear viscosity ($\eta$0).

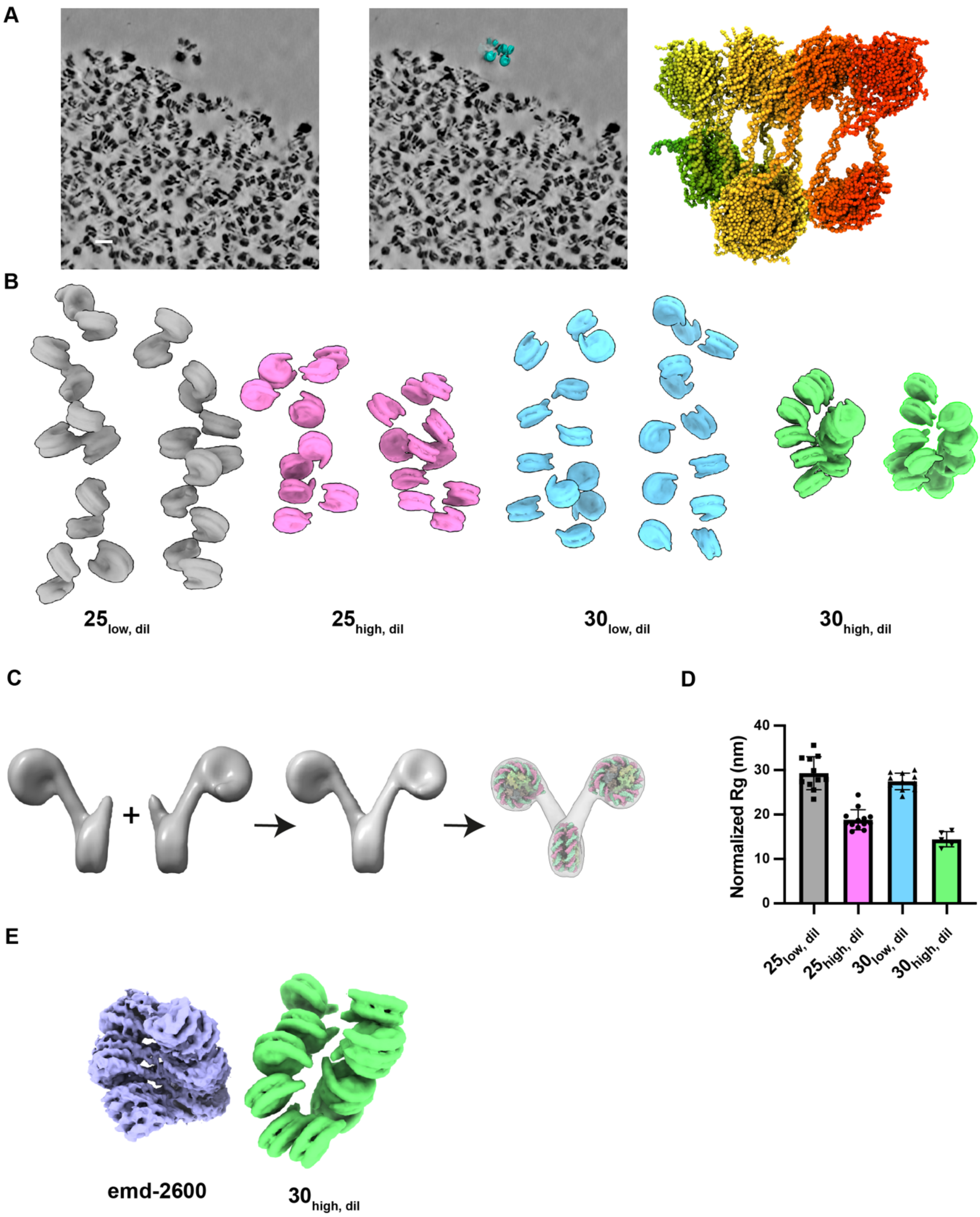


**Fig. S1. Tracing and quantitative analyses of nucleosome arrays in the dilute phase.**

**(A)** Schematic depiction of the computational approach used to reconstruct the missing histone tails and increase the resolution of a single cryo-ET traced chromatin array at high salt conditions. The first two panels illustrate: first an individual chromatin array traced from cryo-ET data outside of the condensate, and then the result of coloring the nucleosomes according to their sequential nucleosome number. The third panel shows the result of reconstructing the cryo-ET fiber computationally.

**(B)** Representative nucleosome arrays under the specified conditions, 25 bp chromatin in low-salt dilute ($25_{low, dil}$) and high-salt dilute ($25_{high, dil}$) phases, and 30 bp chromatin in low-salt dilute ($30_{low, dil}$) and high-salt dilute ($30_{high, dil}$) phases.

**(C)** Modelling of the tri-nucleosome models for visualization. Two identical subtomogram-averaged di-nucleosome structures were aligned at the central nucleosome to generate a tri-nucleosome configuration. Mononucleosome structures (PDB: 6pwe) were fitted into the nucleosome density to enhance visualization.

**(D)** Statistical characterization of $R_g$ measurements (normalized to 12 nucleosome length, see Methods) for traced 25 bp and 30 bp chromatin in low salt and high salt conditions. Error bars represent the standard deviation of the measurements for $\geq 5$ arrays.

**(E)** Comparison of the 11 Å resolution single-particle cryo-EM map of a 30 bp linker length chromatin fiber in the presence of linker histone H1.4 (emd-2600) with one of the traced 30 bp chromatin fibers in the high salt dilute phase.

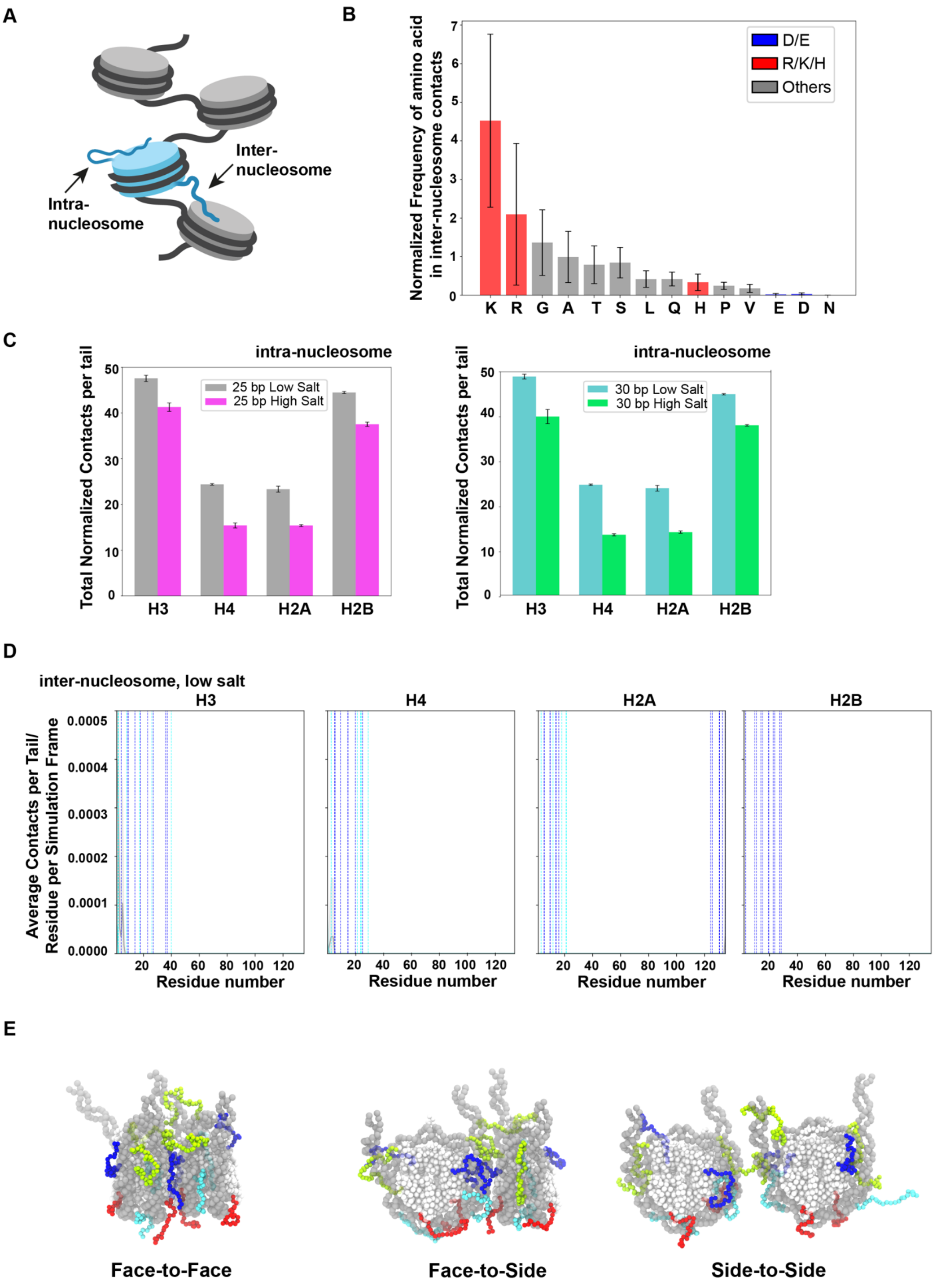


**Fig. S2. Simulation of histone tails in the dilute phase.**
**(A)** Schematic illustration of histone tail-mediate intra-nucleosome and inter-nucleosome interactions.
**(B)** Normalized frequency of amino acid pair inter-nucleosome contacts involving different residue types.
**(C)** Total number of contacts mediated by the N-terminal histone tails (H3, H4, H2A(N), and H2B) of one nucleosome and the DNA (nucleosomal and linker) and histones (core ) of the same nucleosome (intra-nucleosome nucleosome). The contacts are computed from molecular dynamics simulations of various different (see Methods) chromatin arrays where nucleosomes have been restrained to maintain their cryo-ET positions. The number of contacts is defined as the total number of amino acid–phosphate and amino acid–amino acid pairs closer than a set cutoff distance (the cutoff value depends on the identity of the interacting particles as described in Methods), per tail type per nucleosome averaged amongst the different arrays per condition and across the simulation trajectory. The error bars represent one standard deviation from the mean for the averaged totals. The plots compare the results for 25-bp linker chromatin arrays (left plot) and 30-bp chromatin arrays (right plot) under both high and low salt conditions.
**(D)** Number of tail-mediated inter-nucleosome contacts in low salt condition. Data for 25 bp fibers is shown in pink and for 30 bp fibers in green. The blue and cyan vertical lines show the positions of lysines and arginines in the tails, respectively. The shading indicates the standard deviation from the mean.
**(E)** Schematic chemically specific coarse-grained representation of tail-mediated inter-nucleosome interactions among nucleosome pairs interacting face-to–face, face-to–side, and side-to–side. DNA beads are shown in yellow, histone core beads in grey, H3 tail beads in green, H4 tail beads in blue, H2A(N) beads in red, and H2B beads in cyan.

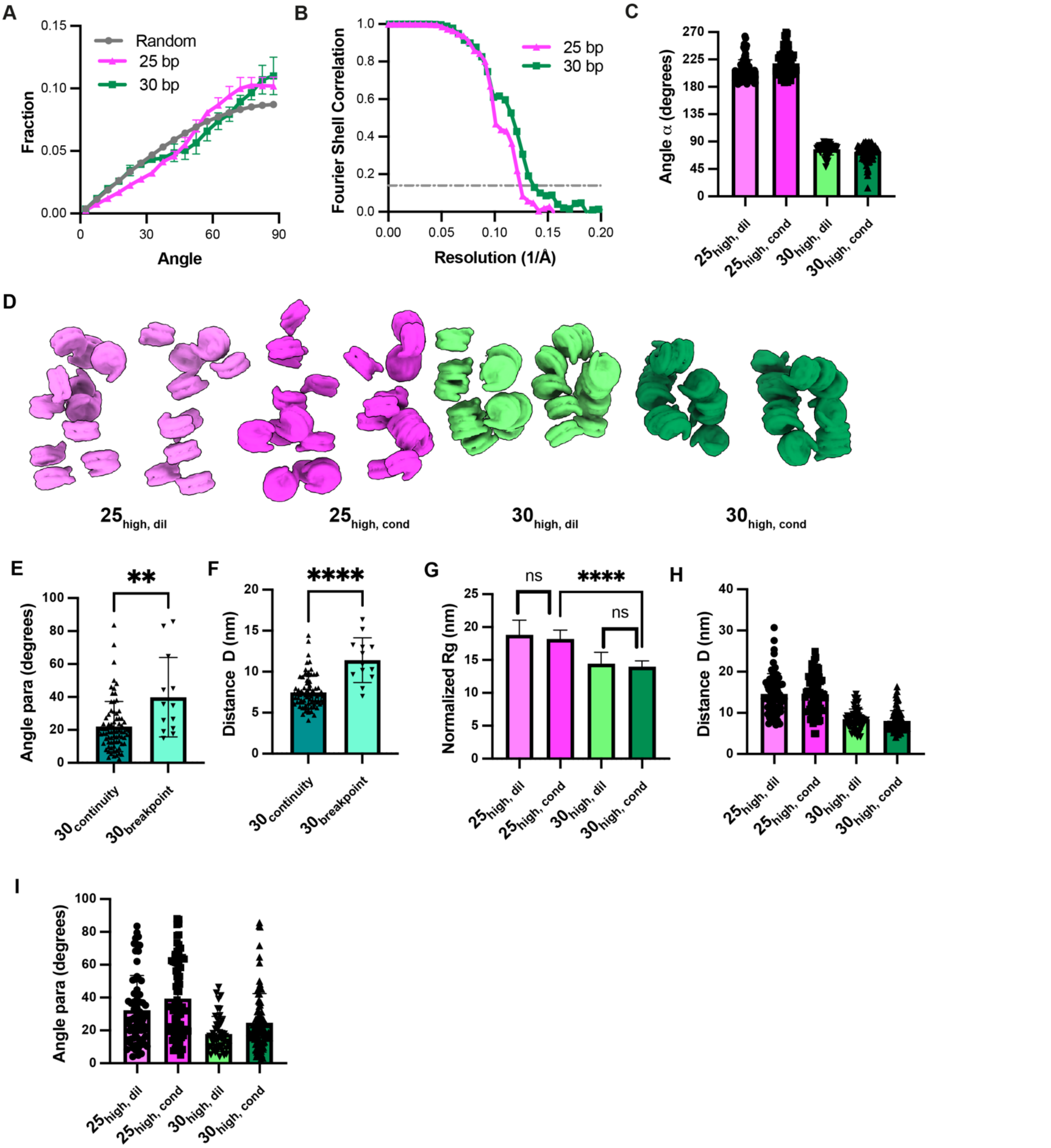


**Fig. S3. Comparison of 25 bp and 30 bp chromatin structures in dilute and condensed phases**

**(A)** Distribution of nucleosome orientations within 25 bp and 30 bp chromatin condensates.

**(B)** Fourier shell correlation from subtomogram averaging of nucleosome structures in condensates, derived from the reconstruction of two half-maps.

**(C)** Comparison of $\alpha$ for 25 bp and 30 bp chromatin under high salt conditions: 25 bp chromatin in the dilute phase with light magenta ($25_{high,\ dil}$) and condensed phase with magenta ($25_{high,\ cond}$), alongside 30 bp chromatin in the dilute phase with light green ($30_{high,\ dil}$) and condensed phase with green ($30_{high,\ dil}$).

**(D)** Structural comparison of nucleosome arrays at the indicated conditions which defined in (C). Each condition includes a pair of 12-mer nucleosome arrays.

**(E-F)** The 30 bp chromatin adopts a 2-start helical structure that typically spans ~4-5 nucleosomes continuously before exhibiting breaks, resembling findings from the single-particle cryo-EM reconstruction of 12-mer arrays with 30 bp spacing (*87*). The continuous regions and breakpoints differ in their distributions of para (panel E) and D (panel F). At the breakpoints, the N and N+2 nucleosomes show increased twisting (higher para) and are more distantly spaced (larger D) compared to the continuous regions.

**(G-I)** Comparison of $R_g$, distance D and dihedral angle para for 25 bp and 30 bp chromatin at the indicated conditions which defined in (C).

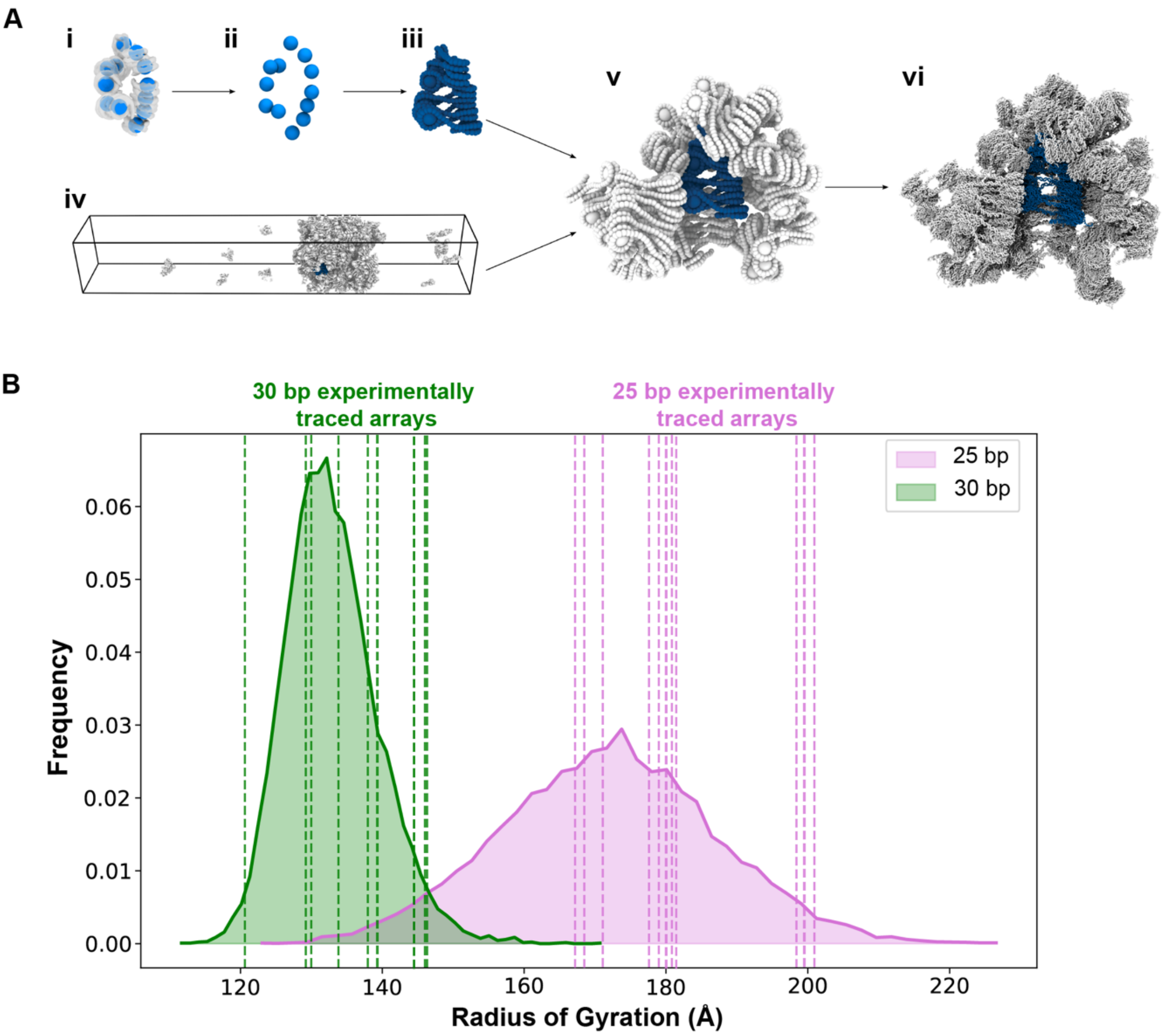


**Fig. S4: Simulations to Probe Histone Tail Interactions inside Chromatin Condensates**

**(A)** Schematic of the multiscale simulation protocol used to enhance the resolution of cryo-ET data for chromatin fibers in condensates down to the amino acid and nucleotide level. Top: Cryo-ET traced chromatin fibers (i) are compared, based on the geometric centers of their nucleosomes (ii), with individual chromatin structures extracted from the simulations (iii). Simulation structures with lower RMSD values (closer matches) to their cryo-ET counterparts are given higher scores. The top-scoring structure from the simulations, highlighted in blue, is selected (iv). These simulations of chromatin condensates are performed using a minimal coarse-grained chromatin model. Panel iv shows part of the slab simulation box with the chromatin condensate coexisting with the dilute phase. Each fiber is represented with one sphere per 5 bp and one ellipsoid for the histone core. Using the top-scoring structure from the condensate simulations, a cluster of neighboring chromatin arrays in contact with the top-scored structure is identified (v, colored grey). This cluster is then backmapped from the minimal coarse-grained model to the chemically specific model, enabling reconstruction to the amino acid and nucleotide level (vi). Finally, molecular dynamics simulations are performed on the high-resolution cluster, applying restraints to preserve the structures of the interacting arrays. This approach allows us to sample the dynamical behavior of histone tails and perform statistical analyses on the inter-nucleosome interactions they mediate.

**(B)** Distribution of chromatin radius of gyration in *in silico* chromatin condensates. Radius of gyration, $R_g$, for chromatin arrays inside simulated condensates of the 25 bp (magenta) and 30 bp (green) chromatin arrays. Each shaded region represents the distribution of $R_g$ values; dashed vertical lines indicate normalized $R_g$ values of experimentally traced arrays in each case. Note that the same coarse grained simulation trajectories used for the analysis here were also reported in (*84*).

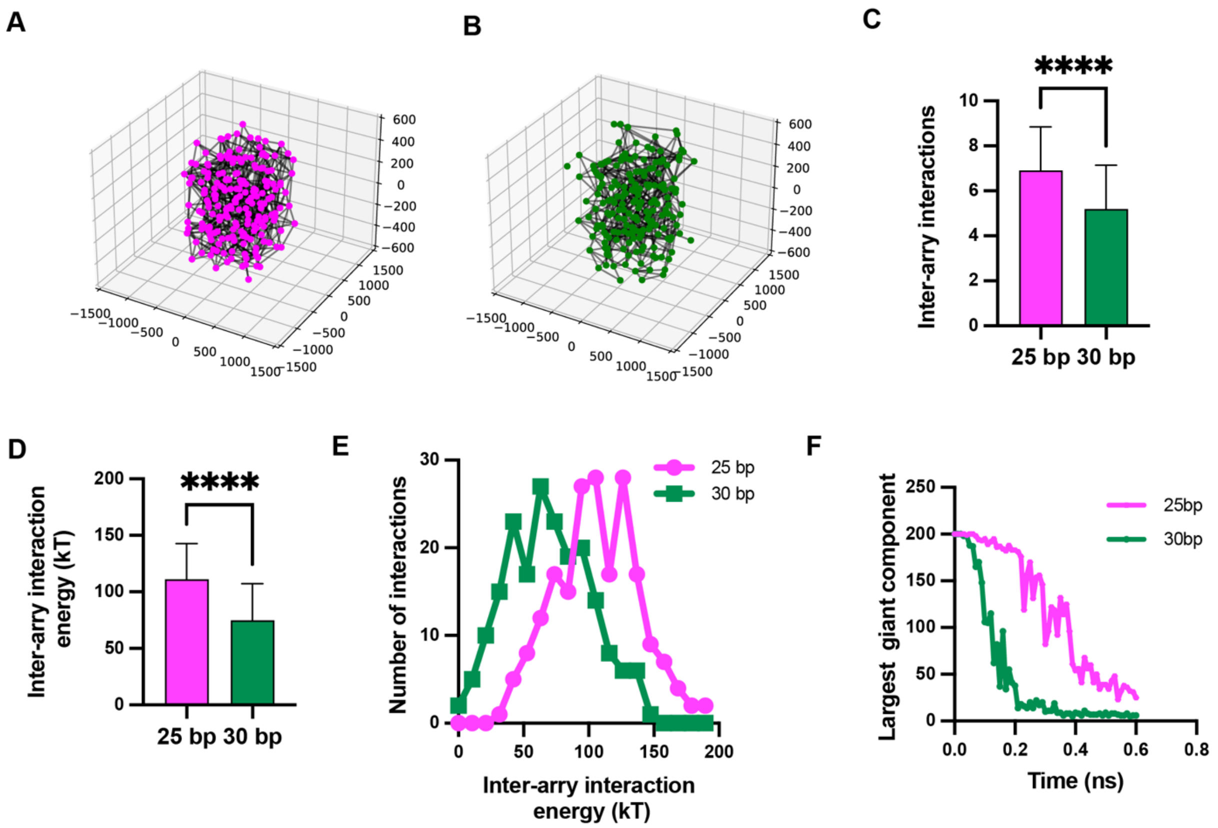


**Fig. S5. Chromatin with 25 bp linkers forms a denser and more connected network compared to chromatin with 30 bp linkers, as shown by coarse-grained simulations.**

**(A-B)** Graph network representations of simulations for 25 bp (A) and 30 bp (B) chromatin. Nodes represent nucleosome arrays, while edges denote interactions between these arrays.

**(C)** The number of inter-array interactions for each nucleosome array in the simulation. Error bars indicate standard deviation of 196 independent measurements.

**(D-E)** The association energy between the nucleosome arrays. In D, error bars indicate standard deviation of 196 independent measurements.

**(F)** The reduction in the size of the largest cluster after deactivating 25% of the interactions between nucleosome arrays.

### Supplementary Tables 1-4

Computational studies (84), structural analysis (148), and single-molecule force spectroscopy (149) have shown that in arrays with linkers longer than ~50 bp, distinctions between 10N and 10N+5 become very small. Such long-linker 10N arrays can sample many different conformations, and are not dominated by stacking (148). Thus, they behave more like 25 bp than 30 bp arrays. In the Supplementary Tables below, we use data on such arrays as a proxy for 25 bp chromatin in our study.

### Supplementary Table 1: Role of histone tails in compacting 25 bp-like chromatin arrays as reported in the literature.

The studies here on long linker arrays are not uniformly consistent with one another, but generally show that all tails have been ascribed importance in array compaction by one or more authors. These results are consistent with our simulations (Fig. 3E-H), where for 25 bp arrays all tails make substantial inter-nucleosome contacts in high salt conditions, due to the configurational heterogeneity of the nucleosomes.

| | Linker length ( repeats*sequence) | Method (salt condition) | Important (Y/N) | Reference |
|---|---|---|---|---|
| H2A | | | | |
| H2B | | | | |
| H2A/H2B | 61 bp (12*5 S rDNA) | Tailless, AUC (2 mM $Mg^{2+}$) | Y | (*150*) |
| | 61 bp (12*5 S rDNA) | Tailless, AUC (0-150 mM NaCl) | N | (*151*) |
| H3 | 61 bp (12*5 S rDNA) | Crosslinking (0-5 mM $Mg^{2+}$) | Y | (*152*) |
| | 61 bp (12*5 S rDNA)<br>61 bp (35*5 S rDNA) | Acetylation mimics, Crosslinking (0-5 mM $Mg^{2+}$) | Y | (*153*) |
| H4 | 61 bp (12*5 S rDNA) | Crosslinking ($Mg^{2+}$) | Y | (*154*) |
| | 50 bp (12*601) | Tailless, Force spectroscopy | N | (*149*) |
| | 61 bp (12*5 S rDNA) | Crosslinking ($Mg^{2+}$) | Y | (*155*) |
| H3/H4 | 61 bp (12*5 S rDNA) | Tailless, AUC (2 mM $Mg^{2+}$) | Y | (*150*) |
| | 61 bp (12*5 S rDNA) | Tailless, AUC (0-150 mM NaCl) | Y | (*151*) |

### Supplementary Table 2: Role of histone tails in compacting 30 bp-like chromatin arrays as reported in the literature.

A number of groups have studied nucleosome arrays with relatively short 10N linkers (30 bp or 20 bp), analogous to the 30 bp array used in our study. These have all found that H4 is important, consistent with large number of inter-nucleosome contacts in our tail simulations (Fig. 3E-H). While other tails have not been extensively examined for short 10N arrays, only H2A has been found to contribute to compaction in the 30 bp system, while H2B and H3 have not. This is again consistent with our observations that after H4, H2A is the tail making the most inter-nucleosome contacts.

| | Linker length ( repeats*sequence) | Method (salt condition) | Important (Y/N) | Reference |
|---|---|---|---|---|
| H2A | 30 bp (12*601) | Tailless, AUC (0.1-1 mM $Mg^{2+}$) | N | (*156*) |
| | 30 bp (12*601) | Mutation, AUC (0.8 mM $Mg^{2+}$) | Y | (*157*) |
| H2B | 30 bp (12*601) | Tailless, AUC (0.1-1 mM $Mg^{2+}$) | N | (*156*) |
| H3 | 30 bp (12*601) | Tailless, AUC (0.1-1 mM $Mg^{2+}$) | N | (*156*) |
| H4 | 30 bp (12*601) | Tailless, AUC (0.1-1 mM $Mg^{2+}$) | Y | (*156*) |
| | 30 bp (48*601) | Cross-linking, EM (1 mM $Mg^{2+}$, 50 mM $K^{+}$) | Y | (*94*) |
| | 30 bp (12*601) | Tailless, crosslinking (0 or 1 mM $Mg^{2+}$) | Y | (*93*) |
| | 30 bp (12*601) | Acetylation mimics, AUC (various salt) | Y | (*158*) |
| | 20 bp (12*601) | Tailless, Force spectroscopy | Y | (*149*) |

**Supplementary Table 3: Role of histone tails in oligomerizing 25 bp-like chromatin arrays as reported in the literature.**

Experimental data show that all tails contribute to chromatin oligomerization for long linkers and a 25 bp linker array, consistent with our simulations, where all tails make substantial intermolecular contacts in the condensed phase.

| | Linker length ( repeats*sequence) | Method (salt condition) | Important (Y/N) | Reference |
|---|---|---|---|---|
| H2A | 61 bp (12*5 S rDNA) | Tailless, Precipitation ($Mg^{2+}$) | Y | (*75*) |
| | 61 bp (12*5 S rDNA)<br>60 bp (12*601) | Acetylation mimics, Precipitation ($Mg^{2+}$) | Y/N | (*159*) |
| H2B | 61 bp (12*5 S rDNA) | Tailless, Precipitation ($Mg^{2+}$) | Y | (*75*) |
| | 61 bp (12*5 S rDNA)<br>60 bp (12*601) | Acetylation mimics, Precipitation ($Mg^{2+}$) | Y | (*159*) |
| H2A/H2B | 61 bp (12*5 S rDNA) | Tailless, Precipitation ($Mg^{2+}$) | Y | (*150*) |
| | 61 bp (12*5 S rDNA) | Tailless, Precipitation ($Mg^{2+}$) | Y | (*151*) |
| H3 | 61 bp (12*5 S rDNA) | Tailless, Precipitation ($Mg^{2+}$) | Y | (*75*) |
| | 61 bp (12*5 S rDNA)<br>60 bp (12*601) | Acetylation mimics, Precipitation ($Mg^{2+}$) | Y | (*159*) |
| | 61 bp (12*5 S rDNA)<br>60 bp (35*5 S rDNA) | Acetylation mimics, Precipitation (0-7 mM $Mg^{2+}$) | Y | (*153*) |
| H4 | 61 bp (12*5 S rDNA) | Tailless, Precipitation ($Mg^{2+}$) | Y | (*75*) |
| | 61 bp (12*5 S rDNA) | Crosslinking, Precipitation (1-6 mM $Mg^{2+}$) | Y | (*160*) |
| | 61 bp (12*5 S rDNA)<br>60 bp (12*601) | Acetylation mimics, Precipitation ($Mg^{2+}$) | Y | (*159*) |
| | 61 bp (12*5 S rDNA)<br>60 bp (35*5 S rDNA) | Acetylation mimics, Precipitation ($Mg^{2+}$) | Y | (*154*) |
| | 25 bp (12*601) | Alanine mutations, Fluorescent imaging (150 mM KOAc) | Y | (*33*) |
| H3/H4 | 61 bp (12*5 S rDNA) | Tailless, Precipitation ($Mg^{2+}$) | Y | (*150*) |
| | 61 bp (12*5 S rDNA) | Tailless, Precipitation ($Mg^{2+}$) | Y | (*151*) |

### Supplementary Table 4: Role of histone tails in oligomerizing 30 bp-like chromatin arrays as reported in the literature.

Experimental data from several groups indicate that histone H4 tail mutations/truncation have the largest inhibitory effect on oligomerization of 30 bp chromatin. This likely arises because the H4 tail is needed both to maintain stacking in the 2-start helix (Supplementary Table 2), and then also for intermolecular interactions when the helix is disrupted, as for arrays in Supplementary Table 3. Thus, the data are not easily interpretable in terms of oligomerization of wild type 30 bp chromatin, nor in comparing to our simulations, where the two-start helix is intact. Data on the other three histone tails is more ambiguous, with truncations in (*156*) showing no effect on chromatin precipitation, but acetylation mimicking K→Q mutations in (*159*) suggesting all tails play roles in precipitation. Our simulations would be more consistent with the latter behaviors, as H3, H2A(N) and H2B all make substantial intermolecular contacts in simulations of the 30 bp condensates.

| | Linker length ( repeats*sequence) | Method (salt condition) | Important (Y/N) | Reference |
|---|---|---|---|---|
| H2A | 30 bp (12*601) | Tailless, Precipitation (0.1-1 mM $Mg^{2+}$) | N | (*156*) |
| | 30 bp (12*601) | Acetylation mimics, Precipitation ($Mg^{2+}$) | Y/N | (*159*) |
| H2B | 30 bp (12*601) | Tailless, Precipitation (0.1-1 mM $Mg^{2+}$) | N | (*156*) |
| | 30 bp (12*601) | Acetylation mimics, Precipitation ($Mg^{2+}$) | Y | (*159*) |
| H3 | 30 bp (12*601) | Tailless, Precipitation (0.1-1 mM $Mg^{2+}$) | N | (*156*) |
| | 30 bp (12*601) | Acetylation mimics, Precipitation ($Mg^{2+}$) | Y | (*159*) |
| H4 | 30 bp (12*601) | Tailless, Precipitation (0.1-1 mM $Mg^{2+}$) | Y | (*156*) |
| | 30 bp (12*601) | Crosslinking, Precipitation (1-6 mM $Mg^{2+}$) | Y | (*160*) |
| | 30 bp (12*601) | Acetylation mimics, Precipitation (various salt) | Y | (*158*) |
| | 30 bp (12*601) | Acetylation mimics, Precipitation ($Mg^{2+}$) | Y | (*159*) |

### Supplementary Table 5: cryo-ET data collection and reconstruction statistics

| | 25 bp chromatin low salt | 30 bp chromatin low salt | 25 bp chromatin condensate | 30 bp chromatin condensate | purified nuclei and cellular chromatin |
|---|---|---|---|---|---|
| Data collection | | | | | |
| Magnification | 33 K | 81 K | 81 K | 81 K | 81 K |
| Voltage (kV) | 300 | 300 | 300 | 300 | 300 |
| Defocus range (µm) | -0.5 | -3 to -4.5 | -3 to -4.5 | -3 to -4.5 | -3 to -4.5 |
| Pixel size (Å) | 2.06 | 1.516 | 1.516 | 1.516 | 1.516 |
| Electron exposure (e–/Å$^2$) | 162 | 120 | 178 | 178 | 178 |
| Tilt-range/step (°) | -60,+60 / 2 | -45,+45 / 3 | -48,+60 / 2 | -48,+60 / 2 | -54,+54 / 3 |
| Tilt-scheme | dose-symmetric, grouping 2 | | dose-symmetric, grouping 3 | | dose-symmetric, grouping 1 |
| Data Processing | | | | | |
| Symmetry imposed | - | - | C1 | C1 | - |
| Initial particle images (no.) | - | - | 109801 | 119577 | - |
| Final particle images (no.) | - | - | 109801 | 119577 | - |
| Map resolution (Å) | - | - | 8.1 | 7.6 | - |
| FSC threshold | - | - | 0.143 | 0.143 | - |

**Movie S1. Overview of multi-scale structural studies on chromatin condensates, spanning histone tail interactions, molecular interactions, and mesoscale condensate properties.**
